# Hijacking of Multiple Phospholipid Biosynthetic Pathways and Induction of Membrane Biogenesis by a Picornaviral 3CD Protein

**DOI:** 10.1101/210708

**Authors:** Sravani Banerjee, David Aponte-Diaz, Calvin Yeager, Suresh D. Sharma, Hyung S. Oh, Qingxia Han, Masato Umeda, Yuji Hara, Robert Y.L. Wang, Craig E. Cameron

## Abstract

RNA viruses induce specialized membranous structures for use in genome replication. These structures are often referred to as replication organelles (ROs). ROs exhibit distinct lipid composition relative to other cellular membranes. In many picornaviruses, phosphatidylinositol-4-phosphate (PI4P) is a marker of the RO. Studies to date indicate that the viral 3A protein hijacks a PI4 kinase to induce PI4P by a mechanism unrelated to the cellular pathway, which requires Golgi-specific brefeldin A-resistance guanine nucleotide exchange factor 1, GBF1, and ADP ribosylation factor 1, Arf1. Here we show that a picornaviral 3CD protein is sufficient to induce synthesis of not only PI4P but also phosphatidylinositol-4,5-bisphosphate (PIP2) and phosphatidylcholine (PC). Synthesis of PI4P requires GBF1 and Arf1. We identified 3CD derivatives: 3CD^m^ and 3C^m^D, that we used to show that distinct domains of 3CD function upstream of GBF1 and downstream of Arf1 activation. These same 3CD derivatives still supported induction of PIP2 and PC, suggesting that pathways and corresponding mechanisms used to induce these phospholipids are distinct. Phospholipid induction by 3CD is localized to the perinuclear membrane, the outcome of which is the proliferation of membranes in this area of the cell. We conclude that a single viral protein can serve as a master regulator of cellular phospholipid and membrane biogenesis, likely by commandeering normal cellular pathways.

**AUTHOR SUMMARY:** Picornaviruses replicate their genomes in association with host membranes. Early during infection, existing membranes are used but remodeled to contain a repertoire of lipids best suited for virus multiplication. Later, new membrane synthesis occurs, which requires biosynthesis of phosphatidylcholine in addition to the other more specialized lipids. We have learned that a single picornaviral protein is able to induce membrane biogenesis and decorate these membranes with some of the specialized lipids induced by the virus. A detailed mechanism of induction has been elucidated for one of these lipids. The ability of a single viral protein to commandeer host pathways that lead to membrane biogenesis was unexpected. This discovery reveals a new target for antiviral therapy with the potential to completely derail all aspects of the viral lifecycle requiring membrane biogenesis.

## INTRODUCTION

Myriad cellular mechanisms exist to thwart viral infection [1–4]. These mechanisms are triggered when a cellular pattern recognition receptor (PRR) engages a virus-associated molecular pattern, for example 5’-triphosphorylated RNA, the absence of 2’-O-methylation of the mRNA cap, double-stranded RNA, among many others [1–4]. PRRs are located at every portal of viral entry into a cell but are particularly abundant in the cytoplasm, the site of replication of most RNA viruses, especially positive-strand RNA viruses. RNA viruses have evolved multiple mechanisms to escape host innate immunity [1–4]. Some mechanisms are specific, for example the use of virus-encoded protein(s) to bind and/or to degrade a PRR [1–4]. One generic approach exploited by positive-strand RNA viruses may be the use of a *replication organelle* for genome replication, which limits surveillance by cellular antiviral defenses [5], although the need to evade host defenses in cell culture may not be absolute [6].

Virus-induced replication organelles, also referred to as replication complexes, are apparent in cells infected by positive-strand RNA viruses within a few hours post-infection [7,8]. Some viruses remodel existing membranes. For example, Flaviviruses (Dengue virus, West Nile virus and Zika virus) induce invaginations of negative curvature in membranes of the endoplasmic reticulum (ER) that appear as vesicle packets or spherules [9]. Alphaviruses (Sindbis virus and chikungunya virus) induce similar structures but use membranes of endosomes or the lysosome instead [10]. In contrast, hepaciviruses (hepatitis C virus, HCV) and picornaviruses (poliovirus, PV; Coxsackievirus B3, CVB3; human rhinovirus; and foot-and-mouth disease virus, FMDV) use organellar or vesicular membranes to induce protrusions of positive curvature that interact to form a distinct, virus-induced entity [11–14].

The creation of sites of genome replication that are only permeable to small molecules creates a dilemma for trafficking of viral proteins to these sites, given the expectation that viral proteins are produced in the cytoplasm. Therefore, production and/or trafficking of viral proteins and formation of the replication organelle need to be coordinated. For years, it was presumed that a combination of interactions between viral proteins and between viral and host proteins would be essential to this coordination [15]. However, several years ago it became clear that the phosphoinositide, phosphatidylinositol-4-phoshate (PI4P), was enriched in the picornavirus and hepacivirus replication organelles [16]. This discovery inspired the hypothesis that PI4P contributes to recruitment of viral and cellular proteins to the replication organelle [16].

Phosphoinositides have a well-established role in cellular protein trafficking and in coupling activation of protein function to phosphoinositide binding [17]. The RNA-dependent RNA polymerases (RdRps) from PV and CVB3 have been reported to bind to PI4P, consistent with this role during infection [16]. PI4P is enriched in the Golgi apparatus (Golgi) [18]. A phosphatidylinositol (PI)-4 kinase (PI4K) produces PI4P from PI. At the Golgi, the type IIIβ and type IIα PI4K are used [18]. PI4KIIIβ is activated by a pathway in which the guanine-nucleotide exchange factor (GEF), Golgi-specific brefeldin A-resistance GEF (GBF1), is recruited to Golgi membrane by an ill-defined mechanism and converts the GDP-bound form of ADP-ribosylation factor 1 (Arf1) to its active GTP-bound form [19]. Arf1 activation is a requirement for activation of PI4KIIIβ; however, the steps between Arf1 and PI4KIIIβ activation are not clear [19]. Further complicating our understanding of steps succeeding Arf1 activation is the fact that numerous Golgi functions are controlled by effectors whose recruitment to membranes also rely on Arf1 activation using ostensibly different and incompletely elaborated mechanisms [20,21].

The mechanism(s) used by viruses to induce PI4P remain a topic of active research. There is good evidence for picornaviruses and hepaciviruses that either PI4KIIIβ or PI4KIIIα can be used [22]. In addition, studies of infected cells have implicated GBF1 and Arf1 as contributors to PI4P induction [16,23]. The most detailed mechanistic studies have been performed using PV and CVB3. The earliest studies concluded that the picornaviral 3A protein recruits PI4KIIIβ to membranes [16,24,25]. However, more recent studies have suggested that PI4KIIIβ recruitment to membranes by 3A uses a mechanism independent of GBF1 and Arf1 [26–29]. These observations have been interpreted to mean that picornaviruses have evolved a mechanism to recruit PI4 kinases to membranes for production of PI4P that is unique relative to that used by the cell.

Here we show that ectopic expression of a single picornaviral protein, the 3CD protein, is sufficient to induce PI4P in cells. PI4P induction appears to reflect the capacity of 3CD to hijack and constitutively activate the normal PI4P biogenesis pathway of the Golgi. GBF1, Arf1-GTP and activated PI4KIIIβ are required for PI4P induction. Amino acid substitutions in the 3D domain of 3CD (3CD^m^) interfere with steps pre-Arf1 activation; those in the 3C domain of 3CD (3C^m^D) interfere with steps post-Arf1 activation. Co-expression of 3CD^m^ and 3C^m^D restore PI4P induction, consistent with 3CD functioning in two discrete steps. The steps targeted by 3CD appear to be those understood least in the cellular pathway. Interestingly, 3CD binds to PI4P-containing membranes but not to membranes containing only phosphatidylcholine (PC). Both 3CD derivatives retain the capacity to bind to PI4P-containing membranes. In addition to PI4P, expression of 3CD also induces phosphatidylinositol-4,5-bisphosphate (PIP2) and PC in cells. Neither 3CD^m^ nor 3C^m^D prevented induction of PIP2 or PC, suggesting distinct mechanisms for induction of these phospholipids relative to PI4P. Lipid induction by 3CD leads to a substantial proliferation of membranes in the perinuclear region of the cell. In summary, we have discovered that a single viral protein is sufficient to commandeer multiple phospholipid biosynthetic pathways and create new membranes for use by the virus in the genesis of its replication organelle. Elucidation of the mechanisms used by 3CD to induce membrane biogenesis may reveal non-genomic mechanisms used by the cell to regulate this process.

## RESULTS

### Expression of PV 3CD Protein in HeLa Cells Induces PI4P Synthesis

Although it is now quite clear that phosphoinositides are exploited by many RNA viruses and mark the sites of genome replication, the viral factor(s) contributing to synthesis of these lipids and corresponding mechanisms are largely unknown. The PV genome encodes a polyprotein that can be divided into structural (P1 in **Fig. 1A**) and non-structural (P2 and P3 in **Fig. 1A**) regions. The P3 region is processed by the protease activity residing in the 3C domain to yield 3AB and 3CD protein as the primary cleavage products (**Fig. 1A**). Several years ago, we reported a PV mutant (GG PV) that expresses 3A and 3BCD as the primary cleavage products[30]. This mutant exhibits a delay in the events that lead up to genome replication, as well as a delayed induction of PI4P biosynthesis [30,31]. This observation implicated 3AB and/or 3CD in the PI4P-induction process.

**Figure 1.**
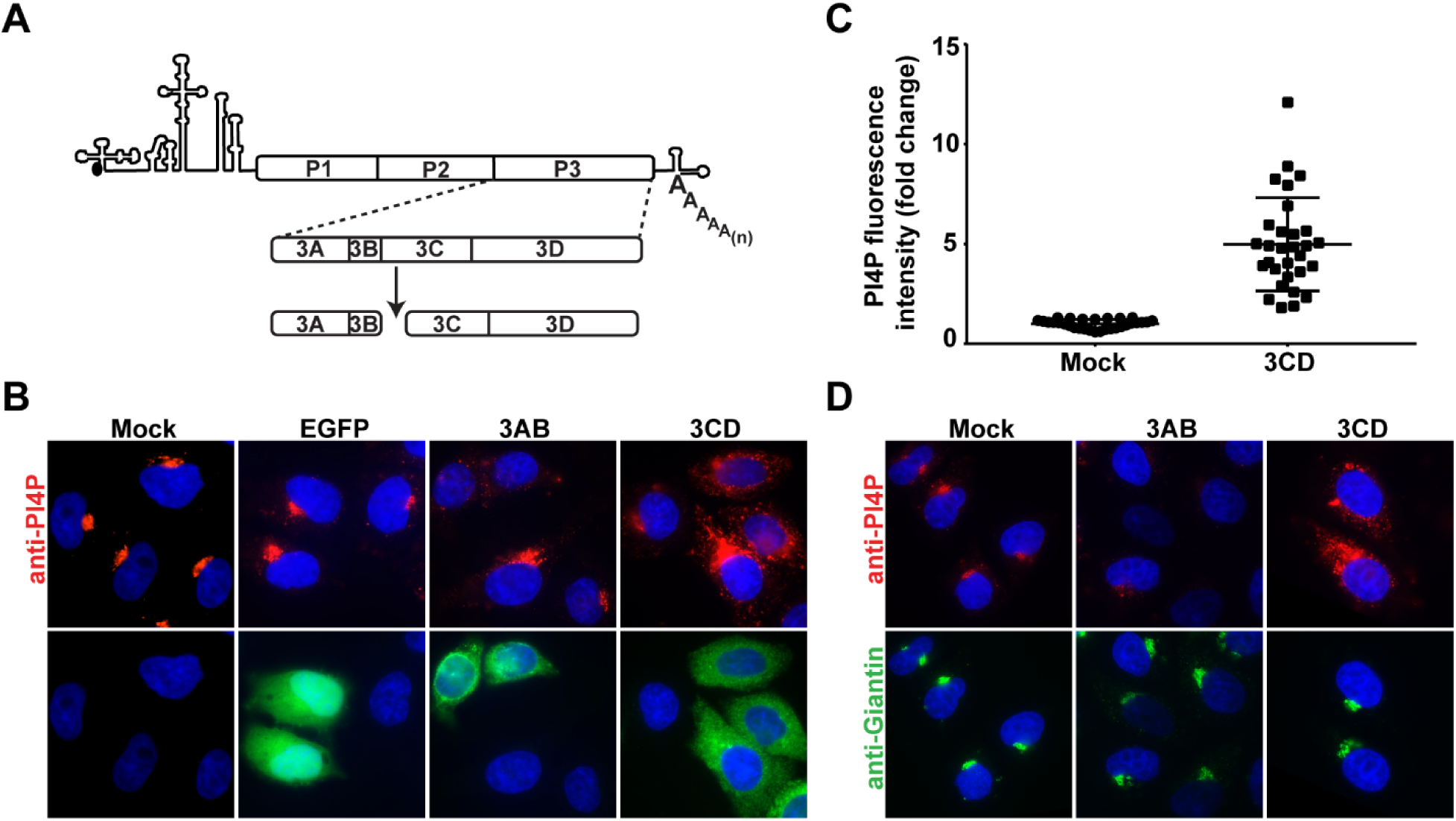
Ectopic expression of poliovirus 3CD causes induction and redistribution of PI4P. **A** Schematic representation of the major pathway for processing of the P3 polyprotein. 3C-encoded protease activity cleaves at the Gln-Gly junction between the 3B and 3C proteins to produce 3AB and 3CD proteins. **(B)** Effect of 3AB and 3CD expression on PI4P levels in HeLa cells. Cells were transfected with in-vitro synthesized mRNA for EGFP, 3AB or 3CD. Four hours post-transfection, cells were immunostained for the presence of PI4P (red) and the expressed protein (green). The nucleus was stained with DAPI (blue). Mock represents cells that were taken through the transfection protocol in the absence of mRNA. Neither the transfection process nor the transfection and translation of EGFP mRNA interfered with levels of PI4P at the Golgi. 3AB diminished PI4P to levels below the limit of detection; 3CD caused both an increase in the level of PI4P as well as its redistribution. **(C)** Quantification of PI4P intensity. The PI4P fluorescence intensity was determined using 11 z-stacks per transfected cell (n=30). The average PI4P intensity of mock-transfected cells was used to normalize the value measured in each cell, the results of which are plotted in the graph. The average of the normalized values were: 1.00 ± 0.04 (SEM) in mock-transfected cells and 4.98 ± 0.43 (SEM) in 3CD-transfected cells. The difference was significant based on a Student’s t-test, which gave a P value of L0.0001. **(D)** Golgi integrity is unaffected by 3AB or 3CD expression. HeLa cells were transfected as above and immunostained for giantin (green) and PI4P (red) 4 h post-transfection. Changes to PI4P caused by expression of 3AB or 3CD do not impact Golgi integrity.

Expression of PV proteins by DNA transfection was problematic, so we used transcription *in vitro* to produce RNA that we capped and polyadenylated *in vitro* to produce mRNAs for 3AB, 3CD and enhanced green fluorescent protein (EGFP), which served as a negative control. The protease activity of 3CD was inactivated by changing the codon for the catalytic Cys residue to one coding for Gly. We transfected these mRNAs into HeLa cells and used immunofluorescence (IF) to monitor protein expression and the fate of PI4P 4 h post-transfection. EGFP expression caused no change in PI4P levels or localization relative to mock-transfected cells (compare EGFP to mock in **Fig. 1B**). 3AB expression caused a loss of PI4P localization at the Golgi (3AB in **Fig. 1B**). 3CD expression caused an increase in PI4P levels, as well as PI4P redistribution (3CD in **Fig. 1B**). PI4P levels increased by an average of five fold (**Fig. 1C**).

PI4P is present at the highest level in the Golgi [18]. PV infection causes Golgi dissolution [32]. One mechanism of Golgi loss is inhibition of GBF1 by 3A protein which interferes with replenishment of Golgi membranes by blocking anterograde transport from ERGIC [24,33,34]. In order to determine if the observed changes to PI4P levels was a consequence of perturbed Golgi integrity, we repeated the experiment described above. We stained cells with an antibody to PI4P to show changes to PI4P and one to giantin to image the Golgi. Expression of neither 3AB nor 3CD impacted Golgi organization (Giantin in **Fig. 1D**).

### Induction of PI4P by PV 3CD Occurs by a Non-genomic Mechanism and Requires GBF1 and PI4KIII

The PV 3CD protein was clearly localized primarily to the cytoplasm (**Fig. 1B**); however, it is known that 3CD can enter the nucleus and alter gene expression [35]. Perturbations to gene expression are thought to require 3C-protease activity, which was inactivated for the purpose of this study. Nevertheless, to formally rule out the possibility that PV 3CD expression perturbs transcription to induce PI4P, we transfected PV 3CD mRNA into cells treated with actinomycin D (AMD). AMD interferes with transcription by all three cellular RNA polymerases [36]. Disruption of RNA polymerase I by AMD causes redistribution of nucleolin in the nucleus [37]; we used this phenomenon as a positive control for AMD treatment (compare anti-Nucleolin -/+ AMD in **Fig. 2A**). The presence of AMD did not interfere with 3CD-mediated induction of PI4P (**Fig. 2A**).

**Figure 2.**
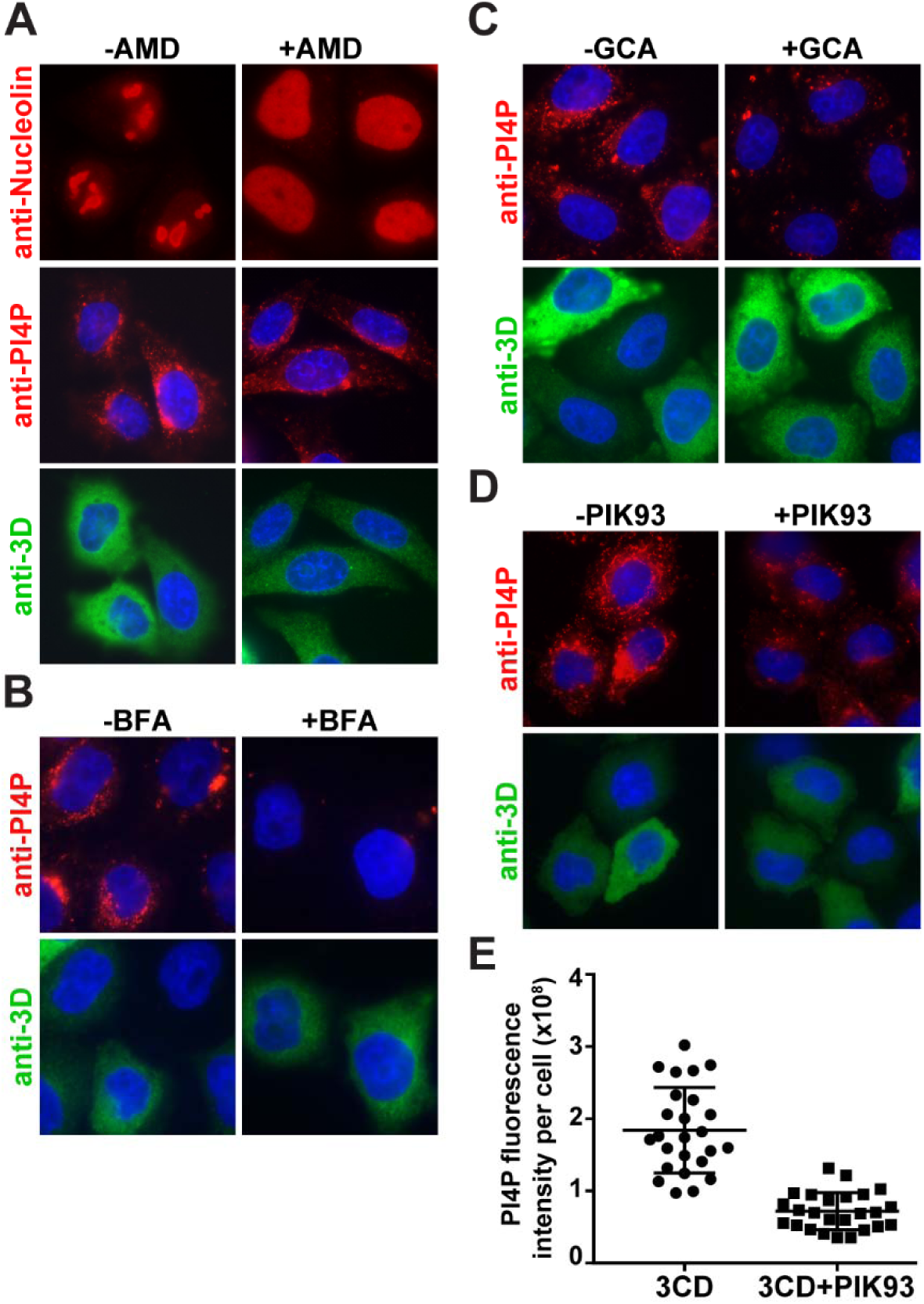
3CD-mediated induction of PI4P does not require host transcription but uses GBF1 and a type III PI4 kinase(s) **A** Effect of actinomycin D (AMD) on 3CD-mediated induction of PI4P. HeLa cells were maintained in the absence or presence of AMD (5 µg/mL) prior to transfection with 3CD mRNA. In one experiment, cells were immunostained for nucleolin (red) to show that AMD treatment was effective. Localization of nucleolin to the nucleolus is lost in the presence of AMD [36]. In a separate experiment, cells were immunostained for PI4P (red) or 3CD (green); the nucleus was stained with DAPI (blue). The presence of AMD did not interfere with induction or redistribution of PI4P. Subsequent experiments were performed as described for AMD but used the following inhibitors: **(B)** brefeldin A (BFA, 2 µg/mL), **(C)** golgicide A (GCA, 10 µM), and **(D)** PIK93 (15 µM). All of these treatments interfered with induction of PI4P. PIK93 was the least effective. **(E)** Quantification of the PI4P levels per cell in the absence (1.8 ± 0.1 × 10^8^) and presence (7.1 ± 0.5 × 10^7^) of PIK93. Error bars represent SEM; Student’s t-test gave a P value of □0.0001.

In order to determine if 3CD-mediated induction of PI4P used aspects of the normal pathway for PI4P production at the Golgi, we measured the impact of brefeldin A (BFA), golgicide A (GCA) and PIK93 on PI4P induction. BFA targets at least three cellular GEFs including GBF1 [38]. GCA only inhibits GBF1 [38]. PIK93 inhibits PI4KIIIβ [39]. 3CD was unable to induce PI4P in the presence of BFA (**Fig. 2B**) or GCA (**Fig. 2C**). The presence of PIK93 reduced the extent to which 3CD was able to induce PI4P (**Fig. 2D**) by more than two fold (**Fig. 2E**).

### Expression of PV 3CD Leads to Arf1 Activation

The preceding experiment suggests that 3CD hijacks the normal cellular pathway, which includes Arf1 activation by converting Arf1-GDP to Arf1-GTP. In order to measure intracellular levels of Arf1-GTP, we used a GGA3 (Golgi-localized, gamma-adaptin ear homology, Arf-binding protein 3) pull-down assay (**Fig. 3A**) [40]. GGA3 is a clathrin adaptor protein that selectively binds to Arf-GTP [40]. We validated the assay using PV-infected cells. Infection caused an increase of Arf1-GTP (**Fig. S1A**) of seven fold (**Fig. S1B**). Expression of 3CD alone caused an increase in Arf1-GTP (**Fig. 3B**) of four fold (**Fig. 3C**).

**Figure 3.**
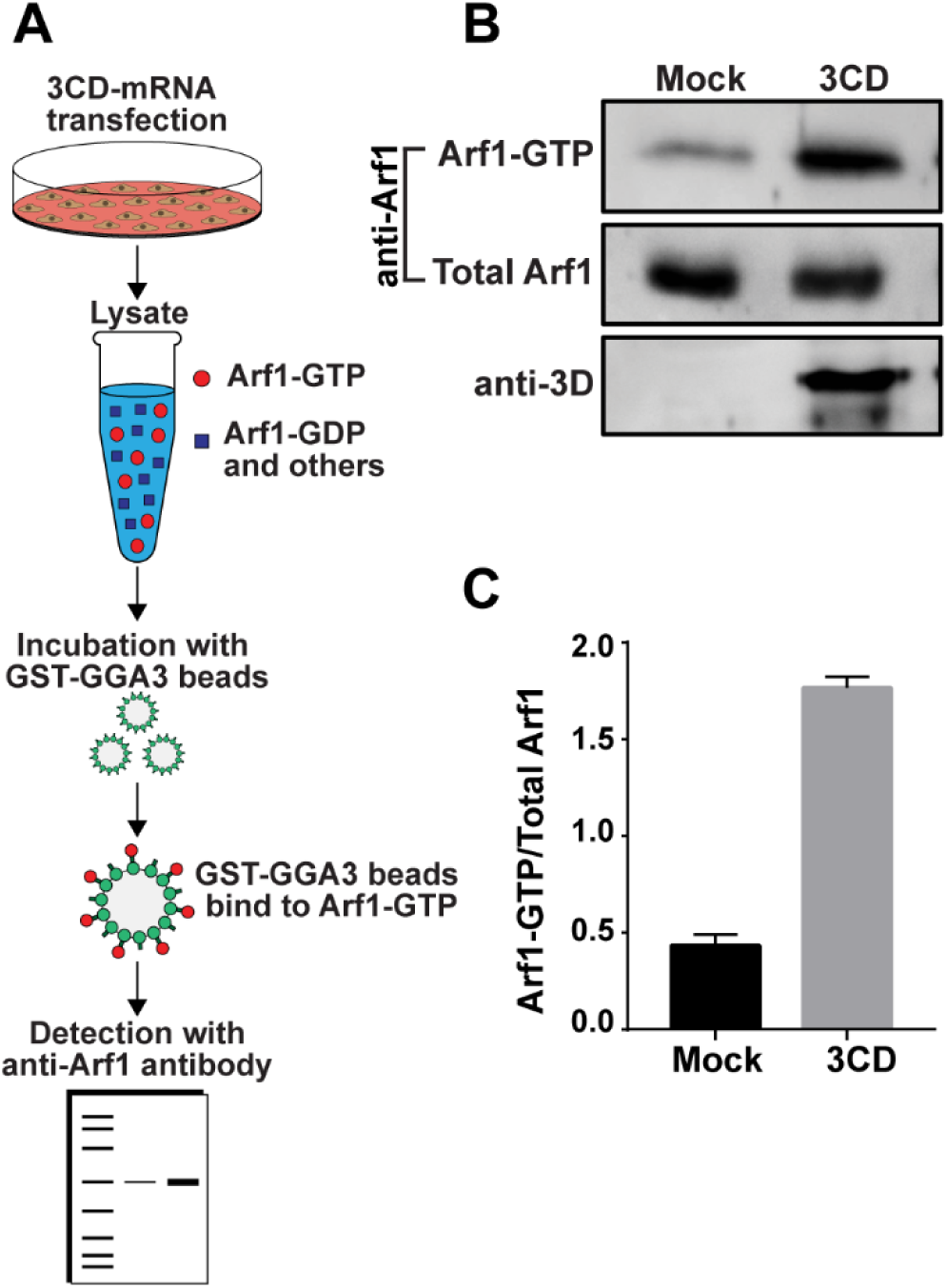
Ectopic expression of 3CD causes activation of Arf1. **A** Schematic of assay used to detect GTP-bound (activated) Arf1. HeLa cells were transfected with or without PV 3CD mRNA. A lysate was prepared using transfected cells and combined with magnetic beads to which the GGA3 adaptor protein was immobilized. Arf1-GTP is uniquely bound by the GGA3 adaptor protein [40]. Beads were isolated, washed, and boiled in SDS-PAGE sample buffer to elute proteins. Eluted proteins were resolved by SDS-PAGE and detected by Western blotting using an anti-Arf1 antibody. **(B)** Detection of Arf1-GTP (bead-eluted material), total Arf1 (from unfractionated lysate), and 3CD (from unfractionated lysate) by Western blot using the indicated antibodies and enhanced chemifluorescence. **(C)** The magnitude of Arf1 activation was determined by quantifying the fluorescence intensity of Arf1-GTP and total Arf1 and reporting the quotient thereof. The average of the values for the quotients (n=3) were: 0.43 ± 0.03 (SEM) in EGFP-transfected cells and 1.77 ± 0.03 (SEM) in 3CD-transfected cells. The difference was significant based on a Student’s t-test, which gave a P value of □0.0001.

### Identification of Genetic Determinants of 3CD Required for PI4P Induction

For years, our laboratory has been interrogating the structure, function and dynamics of PV 3C and 3D proteins, creating a large bank of recombinant PVs with mutations in the corresponding genes. We screened this collection to identify 3CD genes that failed to induce PI4P and identified several. Changing both Leu-630 and Arg-639 in the 3D domain to Asp (referred to throughout as 3CD^m^) interfered with PI4P induction (**Figs. 4A** and **4B**). Changing Lys-12 in the 3C domain to Leu (referred to throughout as 3C^m^D) also interfered with PI4P induction (**Figs. 4A** and **4B**). If each domain functions independently, for example at distinct steps of the PI4P-induction pathway, then each may complement the other. To test this possibility, we co-expressed 3CD^m^ and 3C^m^D. PI4P induction was restored (**Figs. 4A** and **4B**).

**Figure 4.**
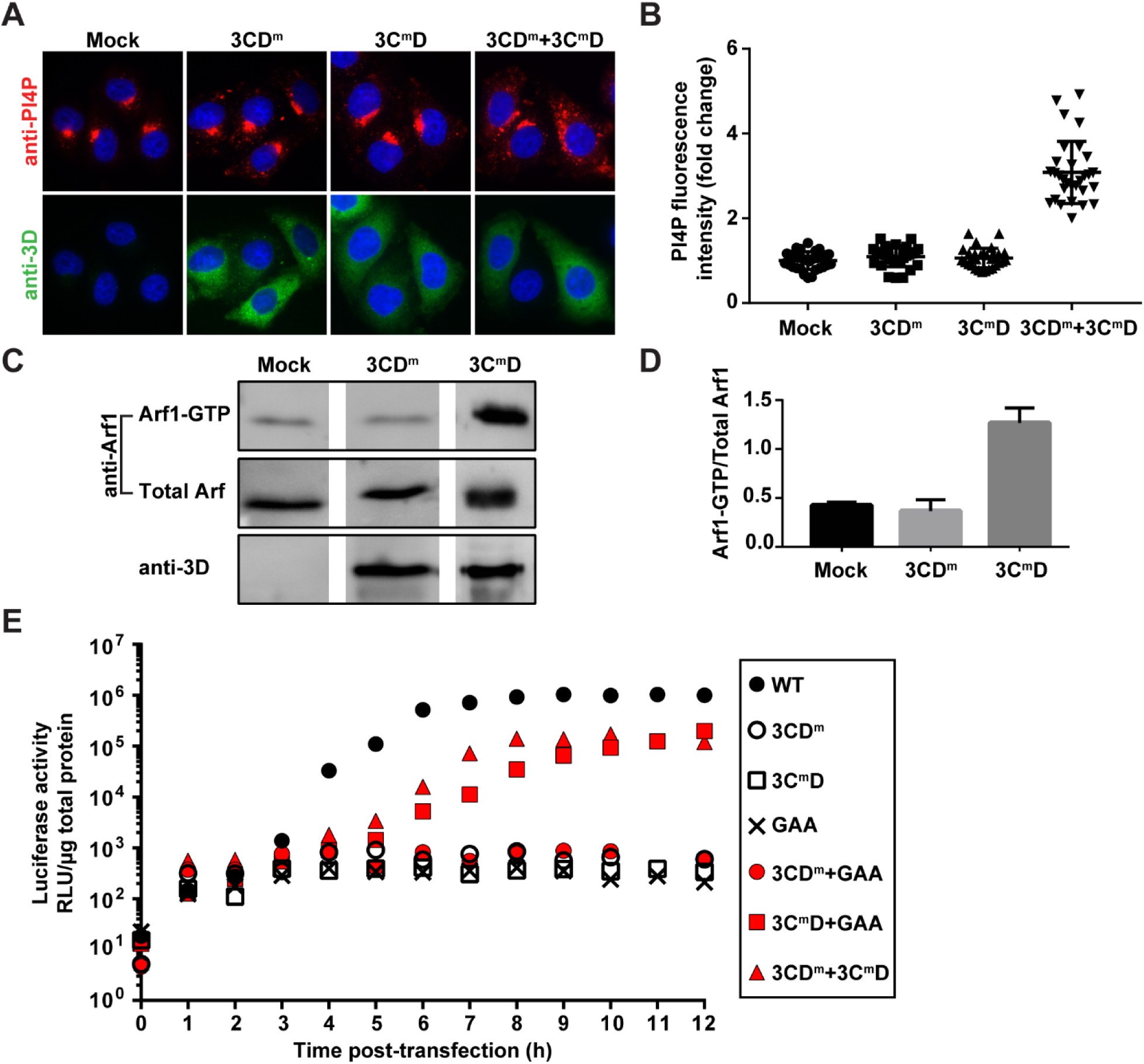
Identification of 3CD variants with substitutions in 3C or 3D domain that impair PI4P induction at distinct steps. **A** 3CD derivatives with substitutions in 3C (3C^m^D) or 3D (3CD^m^) were expressed individually or together in HeLa cells and immunostained for the presence of PI4P (red) and 3CD (green). The nucleus was stained with DAPI (blue). Neither 3CD derivative altered the level of PI4P or its localization; but together the wild-type phenotype was restored. **(B)** Quantification of PI4P intensity per cell was performed as described in the legend to **Fig. 1C**. The average of the normalized values were: 1.00 ± 0.04 (SEM) in mock-transfected cells; 1.06 ± 0.04 (SEM) in 3CD^m^-transfected cells; 1.10 ± 0.04 (SEM) in 3C^m^D-transfected cells; and 3.08 ± 0.13 (SEM) in 3CD^m^ and 3C^m^D co-transfected cells. The level of PI4P induction observed in co-transfected cells was significant when compared to any other experiment based on a Student’s t-test, which gave a P value of L0.0001. **(C)** Activation of Arf1 by the 3CD derivatives was determined as described in the legend to **Fig. 3A**. Only 3CDm failed to induce activation of Arf1. **(D)** The magnitude of Arf1 activation was determined as described in the legend to **Fig. 3B**. The average of the values for the quotients (n=3) were: 0.43 ± 0.02 (SEM) in mock-transfected cells; 0.37 ± 0.07 (SEM) in 3CD^m^-transfected cells; 1.27 ± 0.09 (SEM) in 3C^m^D-transfected cells. For mock vs 3CD^m^, mock vs 3C^m^D and 3CD^m^ vs 3C^m^D, a Student’s t-test yielded P values of 0.4498, 0.0007 and 0.0012, respectively. **(E)** Complementation of 3CD^m^-or 3C^m^D-expressing subgenomic replicons. HeLa cells were transfected with replicon RNA indicated in the key. GAA refers to a replicon expressing a catalytically inactive 3D-encoded polymerase; the corresponding 3CD-GAA should function normally in PI4P induction. Luciferase activity was measured every hour post-transfection as indicated. This experiment was performed twice. Trial one was performed in triplicate; trial two was performed in duplicate. The mean of the values obtained in trial two are shown.

The pathway for PI4P induction is bisected at the Arf1-activation step. Therefore, we used the GGA3 pull-down assay to determine where in the process of PI4P induction each 3CD derivative was impaired. 3CD^m^ failed to activate Arf1, suggesting a defect at a step preceding Arf1 activation (**Figs. 4C** and **4D**). In contrast, 3C^m^D activated Arf1, consistent with a defect at a step succeeding Arf1 activation (**Figs. 4C** and **4D**).

Introduction of 3CD^m^ and 3C^m^D into a PV subgenomic replicon that expresses firefly luciferase completely abolishes genome replication as measured by luciferase activity (**Fig. 4E**). If the observed replication defect is related to functions of 3CD required for formation of the RO, then we should be able to rescue this defect by providing a wild-type copy of 3CD in trans. However, if the defect is related to other genome-replication functions of 3CD, then we should be unable to rescue[41,42]. We delivered the “wild-type” 3CD in the form of a mutant in which the catalytic site of the RdRp has been inactivated by changing the signature GDD sequence to GAA. As expected, the GAA replicon was dead (**Fig. 4E**). Co-transfection of the replicons expressing 3CD^m^ or 3C^m^D and GAA showed complementation of only 3C^m^D, although both the kinetics and yield of luciferase activity were reduced relative to the bona fide wild-type replicon (**Fig. 4E**).

The inability of GAA to complement 3CD^m^ was anticipated because the amino substitutions in the 3D domain of 3CD^m^ interferes with recruitment of the polymerase to the site of genome replication [43]. The combination therefore should be unable to perform genome replication. Co-transfection of 3CD^m^ and 3C^m^D should not have this problem. Each of these two defective replicons complemented the other (**Fig. 4E**), consistent with defects to induction of PI4P biosynthesis contributing to the inability of these replicons to replicate.

While we only identified one 3CD^m^ variant that incapacitated PI4P production, we identified two additional 3C^m^D variants incapable of PI4P induction (**Figs. S2A** and **S2B**). These 3C^m^D variants changed Arg-13 to Leu (referred to as R13L) or Arg-84 to Leu (referred to as R84L). Both variants were capable of Arf1 activation (**Figs. S2C** and **S2D**) and could be complemented to varying degrees by the GAA replicon (**Fig. S2E**). The efficiency with which replication was restored by the GAA replicon could be ordered as follows: R13L > K12L > R84L.

### PV 3CD binds to PI4P-containing Membranes

Recently, we developed an assay to study the interaction of phosphoinositide-binding proteins with membranes using a supported-lipid bilayer prepared in the channels of a microfluidic device [44]. A fluorescence probe in the bilayer senses protein binding, causing fluorescence quenching for the proteins used to date. The system was validated using the Pleckstrin homology (PH) domain from phospholipase C δ1 that binds to PIP2 [44].

The first study demonstrating PI4P induction showed that the 3D domain alone was able to bind to immobilized PI4P [16]. It therefore seemed reasonable to expect that 3CD would also bind to PI4P. To test this possibility formally, we used the chip-based assay to evaluate 3CD binding to PC-based membranes in the absence or presence of PI4P. 3CD bound to the membrane only when PI4P was present (**Fig. 5**). Interestingly, both 3CD^m^ and 3C^m^D retained the capacity to bind to PI4P-containing membranes, suggesting that the ability to be recruited to the appropriate membranes remained intact. A statistically significant difference in the binding affinity for both derivatives relative to wild-type 3CD was noted (**Fig. 5**). 3C^m^D exhibited two-fold higher affinity binding; 3CD^m^ exhibited two-fold lower affinity binding (**Figs. 5B and 5C**). If and how these changes to PI4P-binding affinity correlate to the observed phenotype in cells requires further investigation.

**Figure 5.**
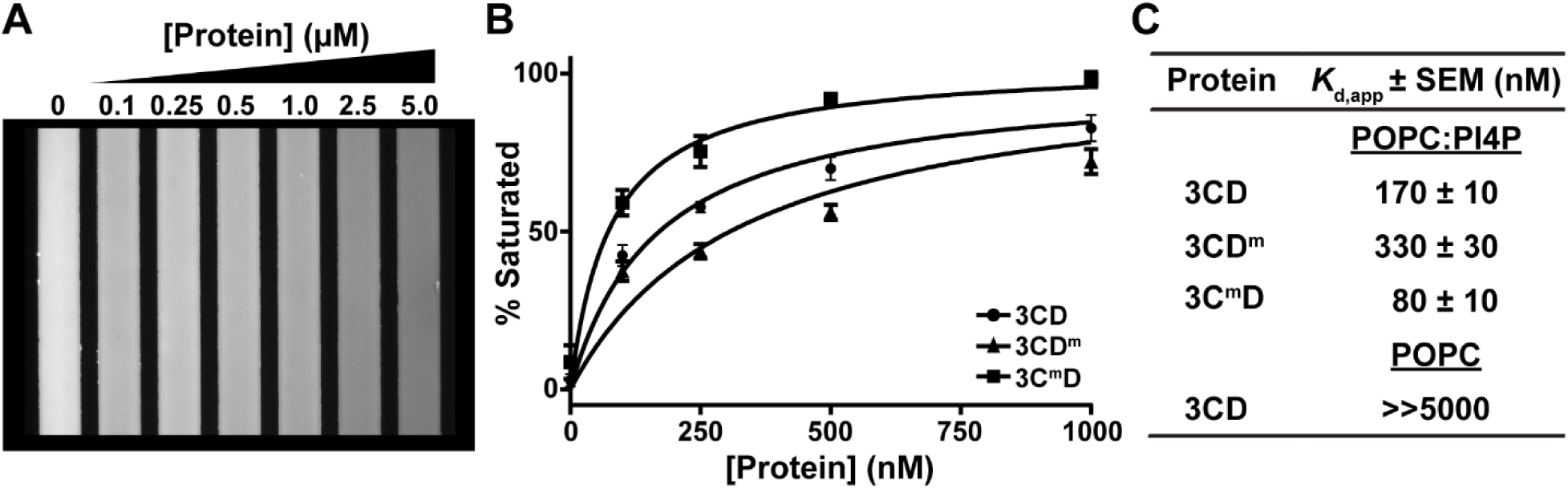
3CD derivatives exhibit modulated binding to PI4P-containing membranes. **A** A microfluidics-based, membrane-binding assay was used to measure membrane binding by 3CD and derivatives thereof [44]. A bilayer is formed in the channels of the microfluidics device. The bilayer contains a phosphatidylcholine (POPC) and a fluorescent derivative of phosphatidylethanolamine (0.5 mol%) in the presence or absence of PI4P (7.5 mol%). Binding of the protein to the membrane quenches fluorescence, which enables measurement of the dissociation constant, *K*_d,app_. **(A)** Image of the fluorescence observed in the channels of the device as a function of 3CD concentration using a 10X objective. **(B)** The change in fluorescence was converted to percent of bilayer saturated (see Materials and Methods) and plotted as a function of protein concentration. Data were fit to a Langmuir binding isotherm with an n=1 to obtain values for *K*_d,app_. Three independent experiments were performed. Error bars represent standard error of the mean. **(C)** Values for *K*_d,app_ measured for 3CD and its derivatives on the indicated bilayer. When using a bilayer consisting of POPC and the fluorescence probe, no change in fluorescence was observed at a concentration as high as 5000 nM.

### Expression of PV 3CD in HeLa Cells Induces Synthesis of PIP2 and PC

Given the completely unexpected capacity of 3CD alone to induce production of PI4P, we decided to evaluate the fate of other phospholipids in PV-infected cells and determine the extent to which 3CD contributes to their production. Immunological reagents were commercially available for the following phosphoinositides: PI3P, PI3,4P2, PI3,5P2 and PI4,5P2 (PIP2). In addition to PI4P, PV infection also induces production of PC [45,46]. We therefore evaluated synthesis of PC here as well. Of the four additional phosphoinositides tested, PIP2 showed the most robust, reproducible difference in PV-infected cells relative to mock-infected cells (**Fig. 6A**). We observed a near 7-fold increase in PIP2 (**Fig. 6A**). As previously reported, PV infection elevated PC levels by 3-fold relative to control (**Fig. 6B**).

**Figure 6.**
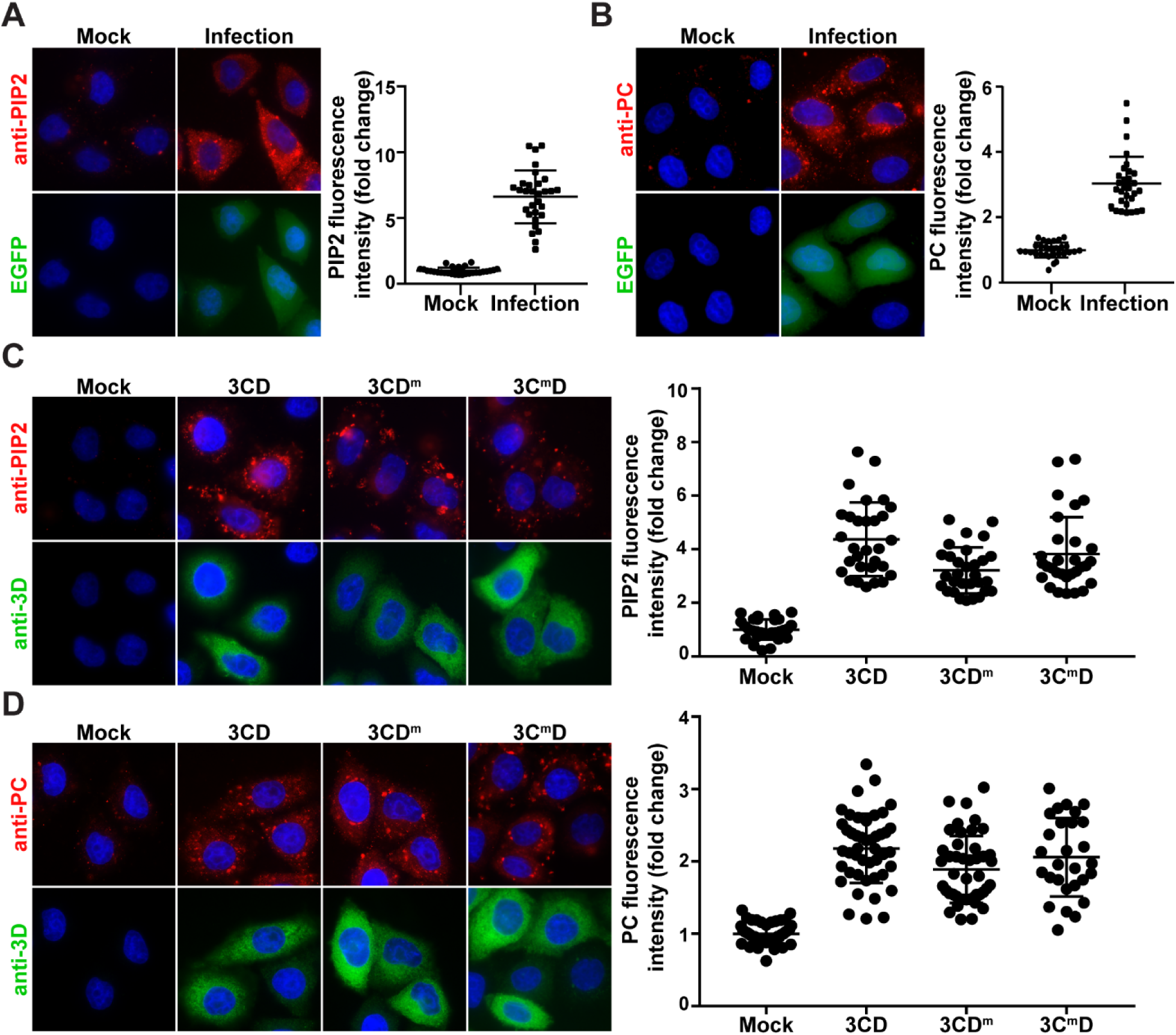
3CD uses distinct mechanisms to induce different lipids. **A** PV infection induces PIP2. HeLa cells were infected with WT PV expressing EGFP at an MOI of 10 and immunostained for PIP2 (red). EGFP (green) indicates infected cells, and DAPI (blue) marks the nucleus. PIP2 was quantified as described for PI4P in the legend to **Fig. 1C**. The average of the normalized values were: 1.00 ± 0.05 (SEM) in mock-infected cells and 6.62 ± 0.37 (SEM) in PV-infected cells. The difference was significant based on a Student’s t-test, which gave a P value of □0.0001. **(B)** PV infection induces PC. Experiment was performed as described in panel A. Cells were immunostained for PC (red). PC was quantified as described for PI4P in the legend to **Fig. 1C**. The average of the normalized values were: 1.00 ± 0.04 (SEM) in mock-infected cells and 3.03 ± 0.15 (SEM) in PV-infected cells. The difference was significant based on a Student’s t-test, which gave a P value of □0.0001. **(C)** Induction of PIP2 by 3CD, 3CD^m^ and 3C^m^D. HeLa cells were transfected with various 3CD mRNAs and fixed 4 h post-transfection. Cells were immunostained for PIP2 (red) and 3CD (green). Nucleus was stained with DAPI (blue). Quantification of PIP2 intensity per cell was performed as described for PI4P in the legend to **Fig. 1C**. The average of the normalized values were: 1.00 ± 0.07 (SEM) in mock-transfected cells; 4.37 ± 0.25 (SEM) in 3CD-transfected cells; 3.21 ± 0.16 (SEM) in 3CD^m^-transfected cells; and 3.82 ± 0.25 (SEM) in 3C^m^D co-transfected cells. The level of PIP2 induction observed in 3CD (WT and mutant)-transfected cells was significant when compared to mock-transfected cells based on a Student’s t-test, which gave a P value of L0.0001. For 3CD vs 3CD^m^, 3CD vs 3C^m^D and 3CD^m^ vs 3C^m^D, a Student’s t-test yielded P values of 0.0002, 0.1264 and 0.0047, respectively. **(D)** Induction of PC by 3CD, 3CD^m^ and 3C^m^D. Experiment was performed as in panel C. Cells were immunostained for PC (red) and 3CD (green). Quantification of PC intensity per cell was performed as described for PI4P in the legend to **Fig. 1C**. The average of the normalized values were: 1.00 ± 0.02 (SEM) in mock-transfected cells; 2.18 ± 0.07 (SEM) in 3CD-transfected cells; 1.89 ± 0.07 (SEM) in 3CD^m^-transfected cells; and 2.06 ± 0.10 (SEM) in 3C^m^D co-transfected cells. The level of PC induction observed in 3CD (WT and mutant)-transfected cells was significant when compared to mock-transfected cells based on a Student’s t-test, which gave a P value of L0.0001. For 3CD vs 3CD^m^, 3CD vs 3C^m^D and 3CD^m^ vs 3C^m^D, a Student’s t-test yielded P values of 0.0041, 0.3129 and 0.1583, respectively.

We evaluated PIP2 and PC levels in 3CD-transfected cells. We observed a 5-fold increase in PIP2 (panel 3CD in **Fig. 6C**) and a 2-fold increase in PC (panel 3CD in **Fig. 6C**) in the presence of 3CD. Like induction of PI4P, 3CD-mediated induction of PIP2 and PC occur by post-transcriptional mechanisms as AMD did not impact levels of these phospholipids (**Figs. S3A** and **S3B**, respectively). Next, we determined if the genetic requirements for PIP2 and PC induction were the same as those for PI4P. 3CD^m^ caused a small but statistically significant (P <0.0001) 1.7-fold decrease in PIP2 levels relative to that observed for wild-type 3CD (compare 3CD^m^ to 3CD in **Fig. 6C**). None of the 3C^m^D variants caused a difference in PIP2 levels relative to wild-type 3CD (**Figs. 6C** and **S3C**). Neither the 3CD^m^ nor the 3C^m^D variants impacted the outcome of 3CD-mediated induction of PC (**Figs 6D** and **S3D**).

### Expression of 3CD in HeLa Cells Induces Membrane Proliferation

Induction of so many different phospholipids by 3CD begs the question: where do they all go? To address this question, we transfected cells with EGFP as a negative control or 3CD mRNA and compared the ultrastructure of the cell as observed by standard transmission electron microscopy. We observed a normal ultrastructure in EGFP-transfected cells (**Fig. 7A**). The Golgi was easily identifiable in the perinuclear region and unremarkable (**Fig. 7B**). In contrast, we observed hypertrophy of membranes in the perinuclear region of 3CD transfected cells (**Fig. 7A**), so much so that the Golgi stacks were no longer distinguishable from the layers of 3CD-induced membranes (**Fig. 7B**). Clearly, Golgi was present and intact based on Giantin staining (**Fig. 1D**).

**Figure 7.**
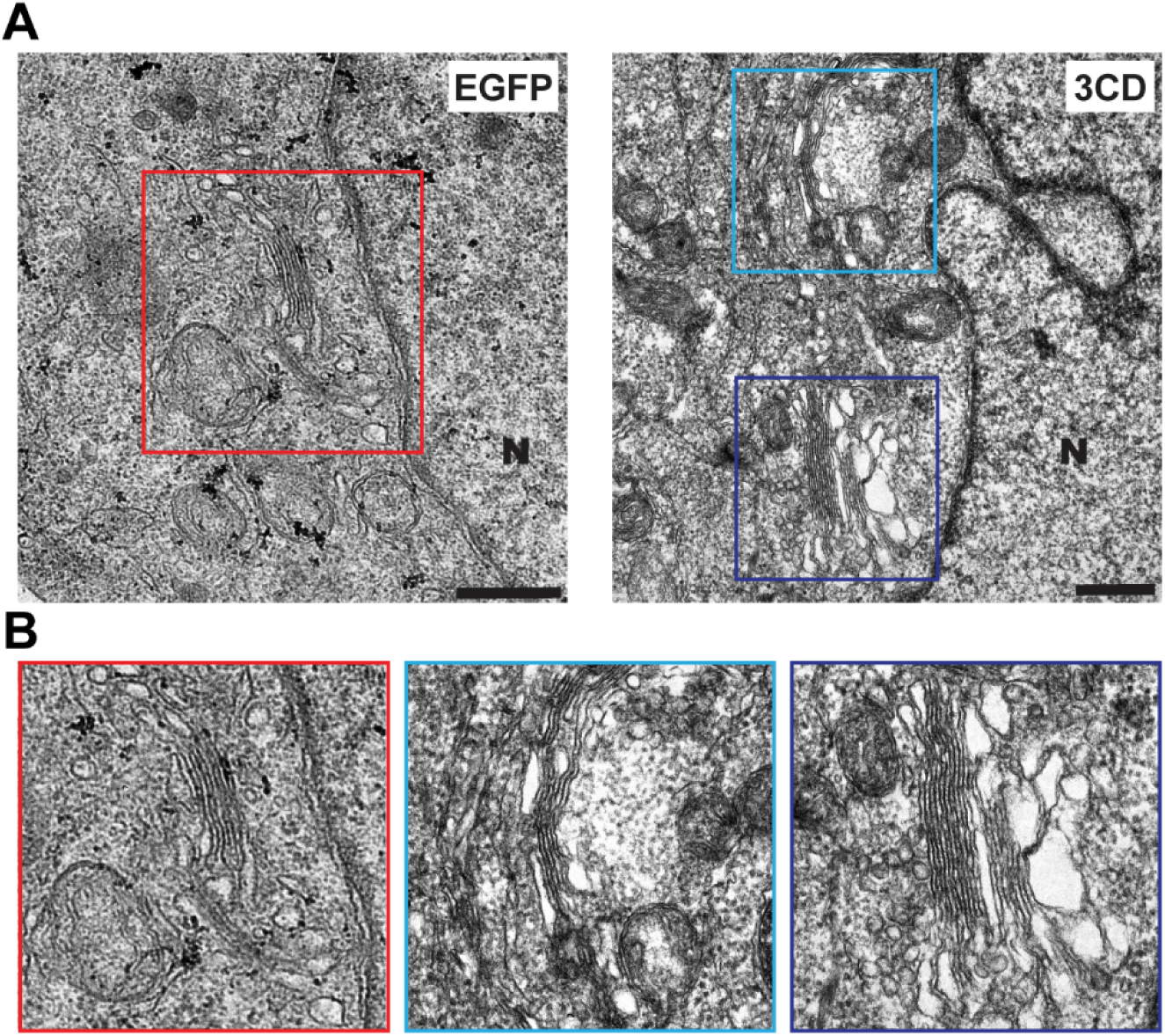
3CD induces membrane biogenesis in the perinuclear region of the cell in the vicinity of the Golgi. eLa cells were transfected with EGFP or 3CD mRNA and then processed for transmission microscopy (TEM) 4 h post-transfection. **(A)** Electron micrograph of a EGFP-transfected cell (left panel) shows the normal ultrastructure of the cell. The nucleus is indicated (N). The Golgi apparatus is located in the perinuclear region of the cell (red box). The comparable view of a 3CD-transfectd cell shows proliferation of membranes in the area normally occupied by the Golgi apparatus (cyan and blue boxes). The Golgi remains intact based on giantin staining (**Fig. 1D**) but is hidden by 3CD-induced membranes. Scale bar indicates 1 µM. **(B)** Magnification of the boxed areas of the perinuclear region of the EGFP-transfected cell (red box) and 3CD-transfected cell (cyan and blue boxes).

## DISCUSSION

Phosphoinositides distinguish one organelle from another, couple activation of protein/enzyme function to appropriate cellular localization and, along with small GTPases and their effectors, enable directional trafficking of proteins and membranes in the cell [17]. In the context of this long-established paradigm of cell biology, the discovery that picornaviruses use PI4P to target proteins to sites of genome replication was logical but also stunning [16]. Since this transformative discovery was made, many laboratories have strived to fill in the gaps between virus entry and formation of the PI4P-rich replication organelle. In the cell, PI4P biogenesis begins with GEF recruitment to the membrane to produce Arf-GTP, which, in turn, recruits effectors that ultimately lead to the recruitment and/or activation of a PI4 kinase [19]. In the specific case studied here, how GBF1 is recruited to membranes and the number and identity of the factors between Arf1-GTP and PI4KIIIβ is unclear (**Fig. 8**).

**Figure 8.**
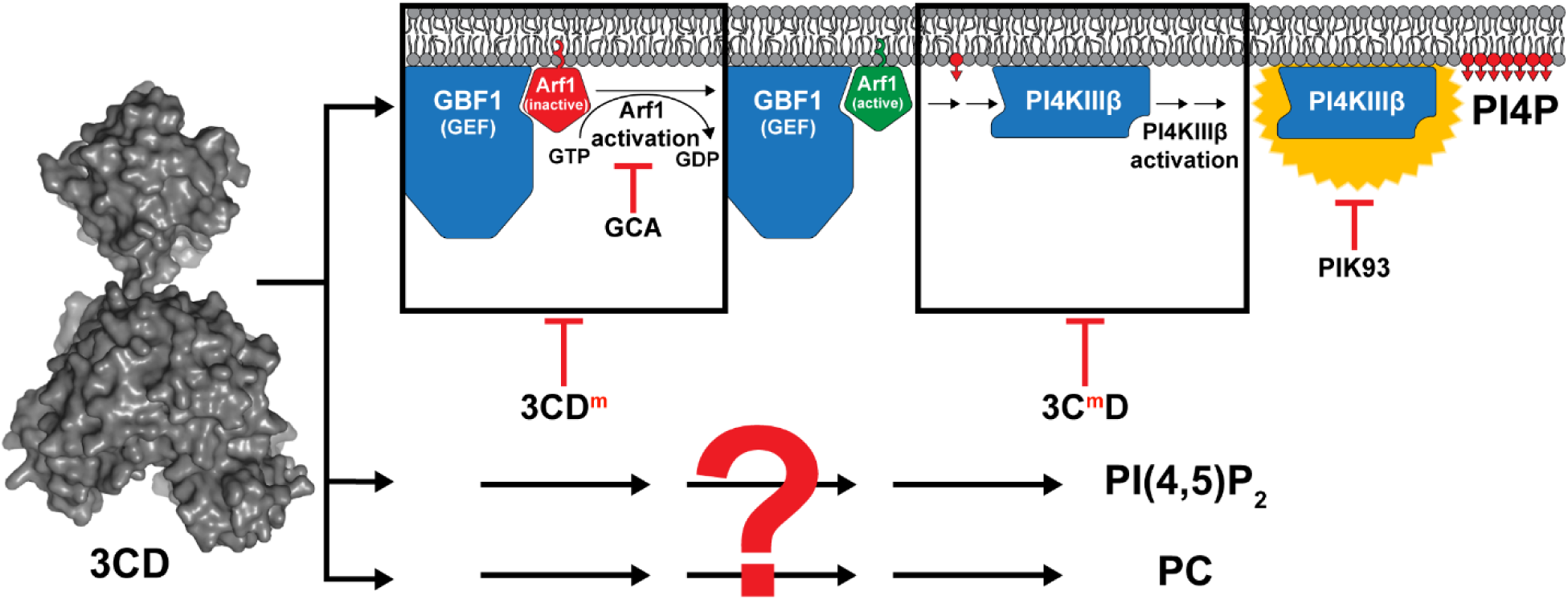
Poliovirus 3CD is sufficient to induce biogenesis of membranes containing PI4P and PIP2. 3CD hijacks the GBF1-Arf1-PI4KIIIβ pathway to induce PI4P biosynthesis. How 3CD recruits GBF1 to the membrane remains unclear. This recruitment event leads to Arf1 activation and is sensitive to golgicide A (GCA). Alteration of the back of the thumb of the 3D domain of 3CD (3CD^m^) interferes with Arf1 activation. Post-Arf1 activation, PI4KIIIβ is recruited to membranes and is itself activated for synthesis of PI4P. PI4P biosynthesis is inhibited by the PI4KIIIβ inhibitor, PIK93. Steps leading to recruitment and/or activation of PI4KIIIβ require the 3C domain of 3CD and are defective in 3C^m^D. The ability of 3CD^m^ and 3C^m^D to complement the other (**Fig. 4**) suggests that 3CD functions minimally at two discrete steps (black boxes). Importantly, these black boxes represent the least understood aspects of the normal pathway of PI4P biosynthesis: recruitment of GBF1 to membranes and recruitment of PI4KIIIβ to membranes and its subsequent activation. Identification of the cellular targets of 3CD may illuminate the cellular proteins participating in these black boxes of the normal cellular pathway of PI4P biogenesis. Biosynthesis of both PIP2 (PI4,5P_2_) and PC is also induced by 3CD. Induction of the synthesis of these lipids occurs by mechanisms distinct from that of PI4P as both 3CD^m^ and 3C^m^D retain the capacity to cause elevation of these lipids (**Fig. 6**). Essentially nothing is known about how 3CD might induce the biosynthesis of these lipids, hence the big red question mark. However, as for PI4P biosynthesis, a non-genomic mechanism of induction is predicted by the inability of actinomycin D to interfere with induction of the biosynthesis of these lipids (**Figs. 2A** and **S3A**).

Conventional wisdom for picornavirology has been that recruitment of the kinase alone or in complex with an unconventional host factor by the picornaviral 3A protein is sufficient to remodel the lipid composition of the replication organelle, formation of which must occur by unrelated mechanisms and directed by other viral factors [16,25–29]. Our study challenges this paradigm by showing that a single viral protein, the 3CD protein, is sufficient to promote membrane biogenesis in the area of the cell between the ER and Golgi, the so-called ER-Golgi intermediate compartment (ERGIC) (**Fig. 7**). These membranes contain the expected PC (**Figs. 6B** and **6D**) and are enriched in PI4P (**Fig. 11**) and PIP2 (**Figs. 6A** and **6C**). Importantly, 3CD-induced changes to levels of PC, PI4P and PIP2 appear to be a deliberate process instead of a transcriptional response to stress (**Figs. 2A** and **S3A,B**).

The mechanism used by 3CD to induce PI4P is the GBF1•Arf1-GTP•PI4KIIIβ pathway (**Figs. 2C-E** and **3**), the normal cellular pathway instead of a virus-contrived pathway. Connections between 3CD and this pathway were made years ago in cell-free systems [47]; however, the field was enthralled by the road less traveled and pursued mechanisms dependent on 3A. Our genetic analysis suggests that 3CD functions both before and after Arf1 activation (**Fig. 4**). The 3D domain contributes to events before Arf1 activation; the 3C domain contributes to events after Arf1 activation (**Fig. 4**). We propose that 3CD hijacks the pathway by infiltrating its two black boxes: (1) recruitment of GBF1 to membranes; (2) recruitment of the effector(s) responsible for PI4KIIIβ activation (**Fig. 8**). Both of these recruitment functions of 3CD would benefit from the ability of 3CD to bind to membranes, an activity reported here to be dependent on PI4P (**Fig. 5**). Identification of the 3CD-interacting factors responsible for PI4P induction may illuminate the mechanism and regulation of the cellular pathway.

The ability of 3AB to promote loss of PI4P from the Golgi (**Fig. 1B**) without interfering with Golgi integrity (**Fig. 1D**) was a surprise. It is known that 3A(B) interacts with GBF1 and the suggestion has been made that the 3A(B)-GBF1 interaction inhibits anterograde transport from ERGIC to Golgi, thus promoting dissolution of the Golgi [24]. For this outcome to occur, transport from ERGIC would have to be impacted instead of transport within the Golgi. What we observe is perhaps GBF1 inhibition by 3AB at the Golgi, the consequence of which may only be inactivation of the normal pathway for maintenance of PI4P levels. Such a mechanism would have the advantage of depleting all PI4P that could misdirect localization of viral proteins, and simultaneously make all GBF1-mediated PI4P synthesis strictly dependent on 3CD. 3AB was much more efficient at clearing PI4P from the Golgi than the PI4KIIIα/β inhibitor, PIK93 (compare PI4P in the 3AB column of **Fig. 1B** to PI4P in the +PIK93 column of **Fig. 2C**). This difference may reflect the fact that PI4P levels at the Golgi are maintained not only by PI4KIIIβ but also by PI4KIIα [18,48]. Indeed, PI4KIIα is responsible for as much as 50% of the PI4P found in cells [48]. This PI4 kinase is not a reported target of PIK93, but our data would suggest that it is a target of 3AB.

Several studies have been published recently reporting that enteroviruses can adapt to growth in the presence of PIK93 [6,49,50]. The suggestion of one group is that induction of PI4P biosynthesis is not absolutely essential for enterovirus genome replication. As discussed above, type III PI4 kinases may not be the only kinases contributing to PI4P biosynthesis as PIK93 is not able to completely abolish induction of 3CD biosynthesis (**Figs. 2D** and **2E**). Resistance to PIK93 maps to 3A-coding sequence, and several alleles have been described [6,49,50]. These substitutions dysregulate processing of 3AB, leading to production of elevated levels of 3A [6,49,50]. Importantly, one group suggests that loss of 3AB correlates with increased levels of PI4P in infected cells in the presence of PIK93 [49]. It is possible that 3A and/or the 3AB derivatives with PIK93-resistance-conferring substitution are unable to interfere with PI4P biosynthesis as well as wild-type 3AB (**Figs. 1B** and **1D**). If this is the case, then resistance to PIK93 does not demonstrate the absence of a need for PI4P but the ability of the virus to shift from utilizing a type III PI4 kinase to another class of PI4 kinases for PI4P biosynthesis.

Both the observation that PV infection induced PIP2 (**Fig. 5A**) and the observation that 3CD alone was sufficient for induction of PIP2 (**Fig. 5C**) were surprises. The existence of plasma membrane-localized pools of PIP2 have been known and studied for a long time [51]. More recently, it has become clear that PIP2 is contained within many intracellular compartments, including ER, Golgi, endosomes and lysosome, and regulates intracellular membrane trafficking, including endolysosomal and autophagic vesicles [51]. Binding of PIP2 to ATG14, a component of the phagophore-inducing PI3 kinase complex, promotes initiation of autophagy [52]. Autophagy contributes substantively to the multiplication of picornaviruses, especially enteroviruses like poliovirus [53–62]. This pathway is thought to be responsible for the non-lytic egress of virus particles. Perhaps 3CD-induced PIP2 during PV infection contributes to formation, function and/or trafficking of the phosphatidylserine-enriched phagophores that engulf PV virions to produce vesicles thought now to be important for viral transmission [63].

Much remains to be learned about PIP2 synthesis during PV infection. PIP2 can be produced from PI4P by a PI4P-5-kinase (PIP5K) or from PI5P by a PI5P-4-kinase (PIP4K) [17]. There are multiple genes for each class; alternative splicing of PIP5K transcripts creates even more isoforms [17]. PIP2-producing kinases are even less well understood than the PI4P-producing kinases [17]. However, PIP2-producing kinases appear to be activated by small GTPases that are regulated by GEFs. Therefore, it is reasonable to propose that the role of 3CD in the mechanism of PIP2 induction will be conceptually similar to that reported here for PI4P induction. 3CD may act upstream of a GEF and downstream of an activated small GTPase to recruit and/or activate the PIP2-producing kinase. The mechanism remains to be defined (**Fig. 8**).

The question of which class of kinases is used will need to be addressed by future experiments. Our observation that 3CD derivatives that fail to induce synthesis of PI4P (**Figs. 4** and **S2**) retain the ability to induce synthesis of PIP2 (**Figs. 6** and **S3**) suggest that 3CD-directed synthesis of PI4P is not required for PIP2. If PI5P is the substrate, then a PIP5K (also referred to as a type II PIPK) would be required. The α and β isoforms of PIP5K reside in the nucleus, and the γ isoform resides in the ER [17,64]. Any or all of these isoforms are feasible candidates in the context of infection because PV infection actually leads to dysregulation of nuclear import and breakdown of the nuclear envelope [65]. In 3CD-transfected cells, it is conceivable that nuclear forms could also be hijacked. 3CD enters the nucleus and could establish sites of PIP2 production at sites of contact between the nucleus and ER [35].

PIP2 is required by other viruses, although there is a paucity of reports of virus-induced synthesis of PIP2. Binding of PIP2 to hepatitis C virus non-structural protein 5A (NS5A) has been shown to promote interactions between NS5A and a host factor required for genome replication [66]. In most other cases, PIP2 is used for some aspect of virion assembly. For example, PIP2 binding to the matrix protein domain of the retroviral Gag polyprotein sends the protein to the plasma membrane as a step in virion assembly [67]. PIP2 binding to the matrix protein (VP40) of Ebola virus is an essential step in virion assembly at the plasma membrane [68]. PIP2 binding to the NP protein of influenza virus in the context of the viral ribonucleoprotein (vRNP) harboring the genome targets the vRNP to the plasma membrane for incorporation into virions [69]. In all of these cases endogenous levels of plasma-membrane associated PIP2 appear sufficient. Use of PIP2 by PV and related viruses for virus assembly fits nicely into this existing paradigm but a mechanism for induction of PIP2 in the cell appears to be required.

In a recent study using representative members of the major supergroups of positive-strand RNA viruses of plants and animals, it was demonstrated that PC is induced by all of the viruses studied, including PV [46]. The mechanism of PC induction used by brome mosaic virus involved recruitment of one PC biosynthetic enzyme and activation of another but details are lacking [46]. Nothing is known about how PV might induce PC. Our inclusion of PC in this study initially was as a negative control. To the best of our knowledge, no pathway using a GEF or a small GTPase is known that leads directly to activation of the enzyme governing the rate-limiting step of PC biosynthesis, CTP: phosphocholine cytidylyltransferase (CCT). Recruitment of CCT to membranes may be sufficient for activation of the enzyme, which could be accomplished by direct interaction with 3CD or a CCT-interacting protein [45]. The observation that 3CD expression caused PC induction motivated us to consider the possibility that phospholipid imbalances resulting from PI4P and/or PIP2 induction triggered a stress response that turned on transcription of genes required for membrane biogenesis. However, the resistance of 3CD-mediated induction of PC to AMD is inconsistent with a transcriptional response (**Fig. S3B**). We suggest the existence of a post-transcriptional or post-translational mechanism that permits cells to respond to an acute need for membrane biogenesis. Perhaps studies of 3CD-mediated induction of PC will illuminate the mechanism.

How is it that a single, viral protein can have so many functions interfacing with host pathways required for phospholipid metabolism and membrane biogenesis, function at the heart of genome replication by binding to all of the cis-acting replication elements for initiation, and also cleave the viral polyprotein to activate viral function and cleave host factors to hijack host ribosomes and antagonize host innate defenses? Our studies of 3CD structure and dynamics have revealed a panoply of conformations achieved by rotating the 3C domain relative to the 3D domain. Each of these conformations have been suggested to expand the proteome and provide unique surfaces for interaction with viral and/or host proteins and nucleic acids [70]. The competency to manipulate the 3CD proteome in a deliberate manner would facilitate elucidation of 3CD-interaction network and uncover mechanisms diluted by the ensemble of states. Understanding the mechanisms used by 3CD to induce these profound changes in the cell will likely illuminate regulatory hubs for membrane biogenesis that have undoubtedly not only been usurped by picornaviruses but other families of RNA viruses as well. Poliovirus and its cousins still hold a lot secrets of mammalian cell biology for future studies to expose.

## MATERIALS AND METHODS

### Cells and culture conditions

HeLa cells (CCL-2) were purchased from American type culture collection (ATCC) and cultured in Dulbecco’s Modified Eagle Medium: Ham’s nutrient mixture F-12 (Gibco) supplemented with 10% heat-inactivated fetal bovine serum (Atlanta Biologics) and 1% Penicillin-Streptomycin (Corning). All experiments were performed at 37 °C in 5% CO_2_.

### Antibodies and other reagents

Antibodies for staining PI4P and PI(4,5)P2 were purchased from Echelon Biosciences. Anti-Giantin and anti-nucleolin were from Abcam and Novus biologicals, respectively. Anti-Arf1 antibody was a generous gift from Dr. Sylvain Bourgoin at Laval University, Canada. Anti-phosphatidylcholine antibody was provided by Dr. Umeda at Tokyo University, Tokyo. Antibodies against PV 3D and 3AB were produced in Cameron lab. All secondary antibodies used for immunofluorescence were purchased from Molecular Probes. Secondary antibody for western blotting was purchased from SeraCare. Inhibitors used in this study including brefeldin A (BFA), golgicide A (GCA), PIK93 and actinomycin D (AMD) were purchased from Sigma-Aldrich.

### Construction of Vectors for Ectopic Expression of Proteins

Plasmids pSB-3AB and pSB-3CD (C147G) used for ectopic expression of proteins, 3AB and 3CD, were constructed by amplifying the gene of interest using pRLuc-RA plasmid (with the protease inactive 3C: C147G) as template and using oligonucleotide pairs mentioned in Table S1. The resulting PCR product was ligated into the pIRES vector (Clonetech) backbone between the EcoRI and NotI unique sites. This caused a loss of the IRES element from the pIRES. To avoid confusion with nomenclature, these cloned plasmids were designated pSB vectors. For making pSB-EGFP, the pEGFP-N1 plasmid was digested with EcoRI and NotI. The resulting EGFP fragment was then ligated into the pIRES backbone between the EcoRI and NotI unique sites. Quickchange mutagenesis was used to construct the mutant 3CD genes carrying point mutations in 3C and 3D using the oligonucleotide pairs mentioned in Table S1. The clones were verified by sending to the sequencing facility at The PSU Genomics Core Facility.

### Engineering Luciferase Subgenomic Replicons with Mutations in 3C

Subgenomic luciferase-encoding replicons harboring the 3C point mutations were constructed by overlap extension PCR using the oligonucleotides listed in Table S1. For both reactions, pRLuc-HpaI-SacII was used as the template [31]. The resulting overlap PCR fragment was digested and ligated in between the HpaI and SacII sites of the pRLuc-HpaI-SacII vector to produce the pRLuc plasmids with specific 3C mutations. The same cloning strategy was followed to construct the other 3C mutant replicons.

### In Vitro Transcription

Linearization of cDNA and transcription was performed as described previously [30]. Briefly, the pSB-and pRLuc-plasmids were linearized with NotI or ApaI and purified with Qiaex II suspension (Qiagen) following the manufacturer’s protocol. RNA was then produced using the linearized plasmid in a 20 µL reaction containing 350 mM HEPES pH 7.5, 32 mM magnesium acetate, 40 mM dithiothreitol (DTT), 2 mM spermidine, 28 mM nucleoside triphosphates (NTPs), 0.025 µg/µL linearized DNA, and 0.025 µg/µL T7 RNA polymerase. The reaction mixture was incubated for 3 h at 37 °C and magnesium pyrophosphate was removed by centrifugation for 2 min. The supernatant was transferred to a new tube and subjected to RQ1 DNase treatment for 30 min at 37 °C. RNA quality was verified by agarose gel (0.8%) electrophoresis.

### RNA Capping and Polyadenylation

The in vitro transcribed pSB-RNAs were purified using the RNeasy kit (Qiagen) using the manufacturer’s protocol and subjected to modification by adding a 5’-end cap and a 3’-end poly(A) tail using the T7 mRNA production kit (Cellscript) following the manufacturer’s protocol. The modified RNAs were further purified using the RNeasy columns and quality was checked by evaluating samples on a 0.8% agarose gel. The addition of poly(A) tail was confirmed by comparing the modified RNA to unmodified RNA.

### mRNA Transfection

2.5 X10^5^ HeLa cells were seeded on coverslips in 6-well plates and transfected 16 h post-seeding. For pSB-mRNAs, 2 µg of column-purified transcripts were transfected using Transmessenger transfection kit (Qiagen) following the manufacturer’s protocol. For pRLuc-RNAs, transfections were performed using 2.5 µg of column purified RNA following the TransIT transfection reagent (Mirus) and corresponding protocol. In all cases, cells were incubated at 37°C and subjected to immunostaining 4 h post-transfection. For studies with inhibitors, transfections were performed in presence of the inhibitors: AMD (5 µg/mL); BFA (2 µg/mL); GCA (10 µM) and PIK93 (15 µM) and the inhibitors were kept on the monolayers for the entire 4 h period.

### Indirect Immunofluorescence

Cells were fixed 4 h post-transfection using 4% formaldehyde in PBS for 20 min followed by washing in PBS and permeabilizing with 20 µM digitonin for 10 min. Digitonin was washed off with three washes of PBS. Cells were then blocked with 3% BSA in PBS for 1 h and incubated with primary antibodies for 1 h. Following washes cells were incubated with the secondary antibodies for 1 h. The processed coverslips were mounted on glass slides using VECTASHIELD Mounting Medium with DAPI (Vector Laboratories). The mounted coverslips were sealed with nail polish. Samples were imaged using Zeiss Axiovert 200 M epifluorescence microscope.

Primary antibodies used were: anti-Giantin (1:400 dilution) for Golgi staining; anti-Nucleolin (1:200 dilution) for nucleolar staining; anti-PI4P (1:200 dilution); anti-PI(4,5)P2 (1:200 dilution); anti-PC (1:40 dilution); anti-3AB (1:400 dilution) and anti-3D (1:100 dilution). All dilutions were made in the blocking buffer.

### Image Processing

For fluorescence intensity quantifications, each image was acquired as a Z-stack file with Z=11 slices.

The sum of intensities was measured using the ZEN 2012 (blue edition) software from Zeiss. The sum fluorescence intensity for each cell was normalized against the average of the sum fluorescence intensities of mock cells.

### Preparing GST-GGA3 Glutathione-Sepharose Beads

BL21-DE3 cells were transformed using a GST-GGA3 expression vector [40]. Transformed cells were plated on NZCYM media containing ampicillin (100ug/ mL) and incubated at 37 °C overnight. BL21-DE3 cells harboring the GST-GGA3 expression vector were then grown overnight in 50 mL NZCYM/Amp at 37 °C. The culture was back-diluted at a 1:25 ratio in 500 mL NZCYM/Amp media and grown at 37 °C until the culture reached 0.4-0.6 OD_600_. Expression of GST-GGA3 was then induced by addition of IPTG to 1 mM and incubated at 37 °C for 6 h. The cell pellet was harvested by centrifugation at 6000 rpm for 10 mins. The pellet (1.8 g) was suspended in 13 mL ice cold bacterial lysis buffer (BLB) containing 50 mM Tris, pH 8.0, 20% sucrose, 10% glycerol, 2 mM DTT, 0.1 mM PMSF and 1 μg/mL each of pepstatin and leupeptin. Cells were lysed using a French press, and cell debris was removed by centrifugation at 25,000 rpm for 30 min at 4 °C. Glutathione-sepharose beads (250 μL) were equilibrated with BLB, by four sequential washes with 1 mL of BLB. Beads were collected after each wash by centrifugation at 400 X g for 2 min at 4 °C. Beads were added to the clarified lysate and incubated at 4 °C for 1 h on a rotating shaker. The beads were then pelleted by centrifugation at 200 X *g* for 30 min and washed six times in 15 mL of BLB. Finally, the beads were suspended in 400 μL of BLB with containing DTT (1 mM). The protein yield was estimated by Bradford assay using BSA as a standard. Beads were aliquoted (40 μg/ tube) for single use and stored at -80 °C.

### Detection of Arf1-GTP in cells using a GST-GGA3 pull-down assay

Activation endogenous Arf1 by transfection of mRNA or by infection was monitored using GST-GGA3 pull-down assay described in [40] with minor changes. Cells were plated at a density of 3 × 10^6^ cells in 100 mm dish 24 h prior to transfection or infection. 3CD (WT or mutant) mRNA (15.5 μg) was transfected into HeLa cells using TransIT-mRNA transfection kit (Mirus). Four hours post-transfection or post-infection cells were lysed at 4 °C in 0.5 mL of lysis buffer (50 mM Tris, pH 7.5, 200 mM NaCl, 10 mM MgCl_2_, 0.1% SDS, 0.5% sodium deoxycholate, 1% triton X-100, 5% glycerol with 0.1 mM PMSF and 1 μg/mL each of pepstatin and leupeptin).

Lysates were incubated on ice for 5 min in presence of 50 μL of CL-4B Sepharose beads (per sample) and then centrifuged for 15 min at 16,000 X *g* and 4 °C. The clarified lysate (500 μg) was incubated with 40 μg of GST-GGA3 bound to Glutathione Sepharose beads for 1 h on a rotating shaker at 4 °C. The beads were then washed three times with 1 mL cold lysis buffer followed by a final wash in 1 mL cold PBS (1X). Liquid was removed beads by quick inversion followed by and a 30 s spin at 6,000 X *g* at 4 °C and removal of residual buffer using a pipetman. Bound proteins were eluted from beads by adding 30 μL of 2X SDS-PAGE sample buffer and incubating at 65 °C for 10 min. Beads were pelleted by centrifugation at 6,000 X *g* for 2 min at room temperature. Twenty μL of the supernatant was resolved by SDS polyacrylamide gel electrophoresis. Endogenous, activated Arf1 was detected using anti-Arf1 antibody. For Total Arf1 was detected using the same antibody, but the sample was prepared from 10 μg of the clarified lysate. Alkaline phosphatase activity associated with the secondary antibody was detected using the ECF reagent (GE Healthcare) and quantified by imaging the fluorescence signal using the Syngene G-box.

### Luciferase Activity Assay

Subgenomic luciferase assays were performed as described previously [30] with the following modifications. Subgenomic replicon RNA (5 µg of in vitro transcribed RNA) was electroporated into HeLa cells. The cells were incubated in normal growth media (DMEM/F12 supplemented with 10% fetal bovine serum, 1% penicillin/streptomycin, 5 mL/1 × 10^6^ cells) and 1 × 10^5^ cells were harvested and lysed using 100 µL of 1X cell culture lysis reagent (CCLR, Promega) at the indicated times post-electroporation. Luciferase activity was measured by adding equal volume of firefly luciferase assay substrate (Promega) to cell lysate and the reaction mixture was applied to a Junior LB 9509 luminometer (Berthold Technologies) to read relative light units (RLU) for 10 s. Relative light units (RLU) were then normalized based on the total protein concentration determined by Biorad protein assay reagent (BioRad) for each sample.

### Transmission electron microscopy

HeLa cells were transfected with mRNAs expressing PV-3CD or EGFP and 4 h post-transfection cells were fixed and embedded for TEM studies as described previously [31]. Briefly, cells were harvested and fixed with 1% glutaraldehyde, washed with 0.1 M cacodylate (sodium dimethyl arsenate, Electron Microscopy Sciences) twice for 5 min each, incubated in 1% reduced osmium tetroxide containing 1% potassium ferricyanide in 0.1 M cacodylate for 60 min in the dark with one exchange and washed two times with 0.1 M cacodylate again. En bloc staining was performed with 3% uranyl acetate in 50% ethanol for 60 min in the dark. Dehydration was carried out with different concentrations of ethanol (50, 70, 95 and 100% for 5-10 min) and 100% acetonitrile. Embedding was performed overnight with 100% Epon at 65 °C. The embedded sample was sectioned with a diamond knife (DiATOME) to slice 60-90 nm thickness by using ultra microtome (Reichart-Jung). The sectioned sample was placed on copper grid (Electron Microscopy Sciences), stained with 2% uranyl acetate in 50% ethanol followed by lead citrate staining for 12 min. The grid was washed with water and the grid was dried completely. The image was obtained using FEI Tecnai G2 Spirit BioTwin located in The PSU Electron Microscopy Facility.

### Supported Lipid Bilayer (SLB) Binding Experiments

A microfluidic platform was employed to test the interaction between the viral protein and 7.5 mol% PI4P SLBs. SLBs containing 99.5 mol% POPC and 0.5 mol% oSRB-POPE, a pH-sensitive fluorescent probe conjugated to a lipid, served as a negative control. Synthesis of oSRB-POPE was described previously [44]. A PDMS block was fused to a borosilicate glass by use of a plasma-oxygen treatment. The fabrication of the PDMS device was described previously [44]. SLBs were formed on the borosilicate glass of each micro channel by spontaneous fusion and rupturing of the SUVs after treatment with 0.2 N HCl [44]. Running buffer (20 mM HEPES, 100 mM NaCl, pH 7.0) was flowed through each channel for 30 min remove excess vesicles. In order to equilibrate the SLBs to the experimental condition, running buffer containing the appropriate dilution of gel filtration buffer (20 mM HEPES, 20% glycerol, 1 mM BME, 500 mM NaCl, pH 7.0) was flowed through the channels. The fluorescence intensity obtained post-equilibration step served as a reference for each channel. PV-3CD dilutions were prepared using the running buffer. PV-3CD was flowed until the fluorescence intensity in each channel stabilized (30-45 min). Change in fluorescence intensity, normalized to the reference (no protein) channel, was plotted as a function of PV-3CD concentration and then fit to a Langmuir isotherm using the equation below.

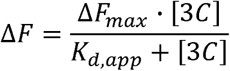

where, *F*_*max*_ represents the normalized fluorescence intensity value at saturation level and *K*_*d,app*_ represents the apparent dissociation constant. Graphs were generated by GraphPad Prism v.6 software. Error bars represent the SEM.

Images were taken with an Axiovert 200M epifluorescence microscope (Carl Zeiss Microscopy) equipped with a AxioCam MRm camera (Carl Zeiss Microscopy) and X-Cite 120 (Excelitas Technologies Corp.) light source was used to take fluorescence images. A 10X air objective was used for imaging along with Alexa 568 filter set (Carl Zeiss Microscopy) with an excitation and emission at 576 nm and 603 nm, respectively. Exposure time was set to 200 msec per exposure with minimal times of exposure throughout experiments. AxioVision LE64 v.4.9.1.0 software (Carl Zeiss Microscopy) was used to process the images.

### *Expression and Purification of* 3C^m^D and 3CD^m^

pSUMO plasmids encoding 3CD^m^ or 3C^m^D was transformed into Rosetta (DE3) competent cells [71]. Cells were grown to an OD of 1.0, harvested by centrifugation (5,400 x g, 10 min, 4 °C), and washed with 30 mL of buffer (10 mM Tris and 1 mM EDTA, pH 8.0) per liter of bacterial culture, and re-centrifuged. Each gram of cell pellet was suspended in 5 mL of lysis buffer (20 mM HEPES, 10% glycerol, 5 mM imidazole, 10 mM β-mercaptoethanol [BME], 500 mM NaCl, 1 mM EDTA, 1.4 μg/mL pepstatin A, 1.0 μg/mL leupeptin, pH 7.0). Suspended cells were homogenized by a Dounce homogenizer and lysed by two passes through a French press at a pressure of 1,000 psi. Phenylmethylsulfonyl fluoride (PMSF) was added to the cell lysate at a final concentration of 1 mM. The suspension was clarified by centrifugation (74,000 x g, 30 min, 4 °C). The supernatant was loaded onto a Nickel-nitrilotriacetic (Ni-NTA) resin (Thermo Fisher Scientific Inc.), which was previously equilibrated with 10 column volumes (CV) of equilibration buffer containing 5 mM imidazole at 1 mL/min using a peristaltic pump. The protein load was passed through the equilibrated Ni-NTA resin at 1 mL/min. To remove contaminants, the loaded resin was washed with 50 CV and 4 CV of equilibration buffer (20 mM HEPES, 20% glycerol, 10 mM β-mercaptoethanol (BME), 500 mM NaCl, pH 7.0) containing 5 mM and 50 mM imidazole, respectively. PV-3CD was eluted into multiple fractions using equilibration buffer containing 500 mM imidazole. Fractions were evaluated by Coomassie stained SDS-PAGE (10.0%) gel. Concentrated fractions were pooled, treated with 1 µg ubiquitin-like-specific-protease 1 (ULP-1) per 1 mg protein of interest, and dialyzed against gel filtration buffer (20 mM HEPES, 20% glycerol, 1 mM BME, 500 mM NaCl, pH 7.0) overnight at 4 °C using a 12-14 kDa MWCO dialysis membrane (Spectrum Laboratories). Dialyzed PV-3CD sample was centrifuged (74,000 x g, 30 min, 4 °C) to ensure non-aggregated protein, and subsequently concentrated to 2 – 5 mL for loading onto a gel filtration column (GE Healthcare). The gel filtration column was run at 1.0 mL of gel filtration buffer per minute, and divided into 34 3-mL fractions. Fractions with the protein of interest were pooled and concentrated with a Vivaspin Turbo 15 (Sartorius, 10 kDa MWCO). The protein concentration was determined by measuring the absorbance at 280 nm (ε_*max*_ = 0.08469 μM^−1^•cm^−1^) with a NanoDrop (ThermoScientific). The ionic strength of the dialyzed protein solution was confirmed with a conductivity meter (Control Company). To assess extraneous protein contamination, dynamic light scattering (DLS) experiments performed with Viscotek 802 DLS instrument at 20 °C. Agarose gel electrophoresis was run to check for any contaminating nucleic acids. Flash frozen aliquots were stored at -80 °C.

### Statistical Analysis

For statistical analysis data were plotted as means ± SEM. The intensity comparisons between control and experimental measurements used an unpaired Student’s t-test in the GraphPad Prism software. The means and P values for pairwise comparisons of all experiments are provided in Table S2, a subset of these are also presented in the appropriate figure legends.

## ACKNOWLEDGEMENTS

We thank Dr. Greg Ning and staff of the electron microscopy core facility for their guidance and assistance. We also thank Dr. Sylvain G. Bourgoin (CHUL, Quebec, Canada) for generously providing the anti-Arf1 antibody.

## SUPPORTING INFORMATION

Figure S1. **Poliovirus infection causes activation of Arf1**.

Figure S2. **Additional 3CD variants with substitutions in the 3C domain exhibit defects to PI4P induction caused by block at a step post-Arf1 activation**.

Figure S3. **Induction of PIP2 and PC biosynthesis by 3CD is not sensitive to actinomycin D and does require induction of PI4P biosynthesis**.

Table S1. **Oligonucleotides used in this study**.

Table S2. **Statistical analysis of all experimental data**.

**Figure S1.**
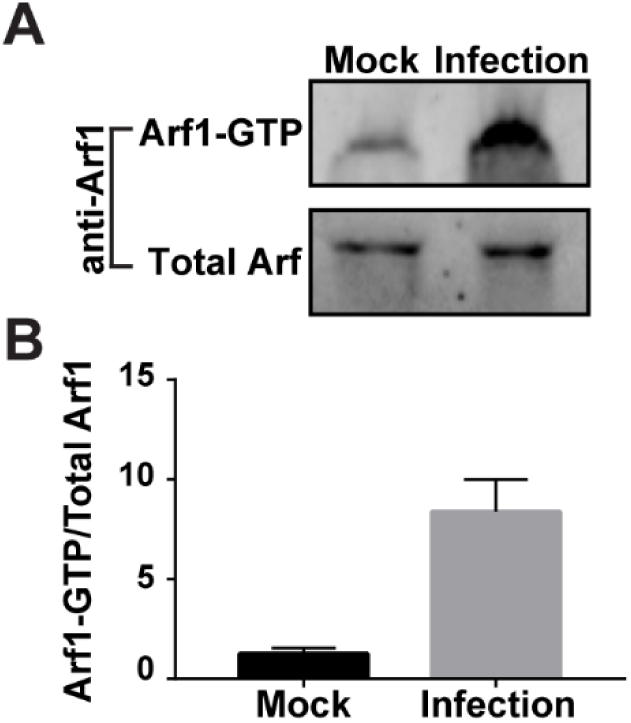
Poliovirus infection causes activation of Arf1. **A** HeLa cells were infected with WT PV at an MOI of 10. A lysate was prepared 4 h post-infection and used for an GST-GGA3-pull-down assay as outlined in **Fig. 3A**. with subsequent western blotting and detection using anti-Arf1 antibody. Arf1-GTP (bead-eluted material) and total Arf1 (from unfractionated lysate) were detected by Western blot using the indicated antibodies and enhanced chemifluorescence. **(B)** The magnitude of Arf1 activation was determined by quantifying the fluorescence intensity of Arf1-GTP and total Arf1 and reporting the quotient thereof. The average of the values for the quotients (n=4) were: 1.27 ± 0.14 (SEM) in mock-infected cells and 8.37 ± 0.03 (SEM) in WT PV-infected cells. The difference was significant based on a Student’s t-test, which gave a P value of 0.0001.

**Figure S2.**
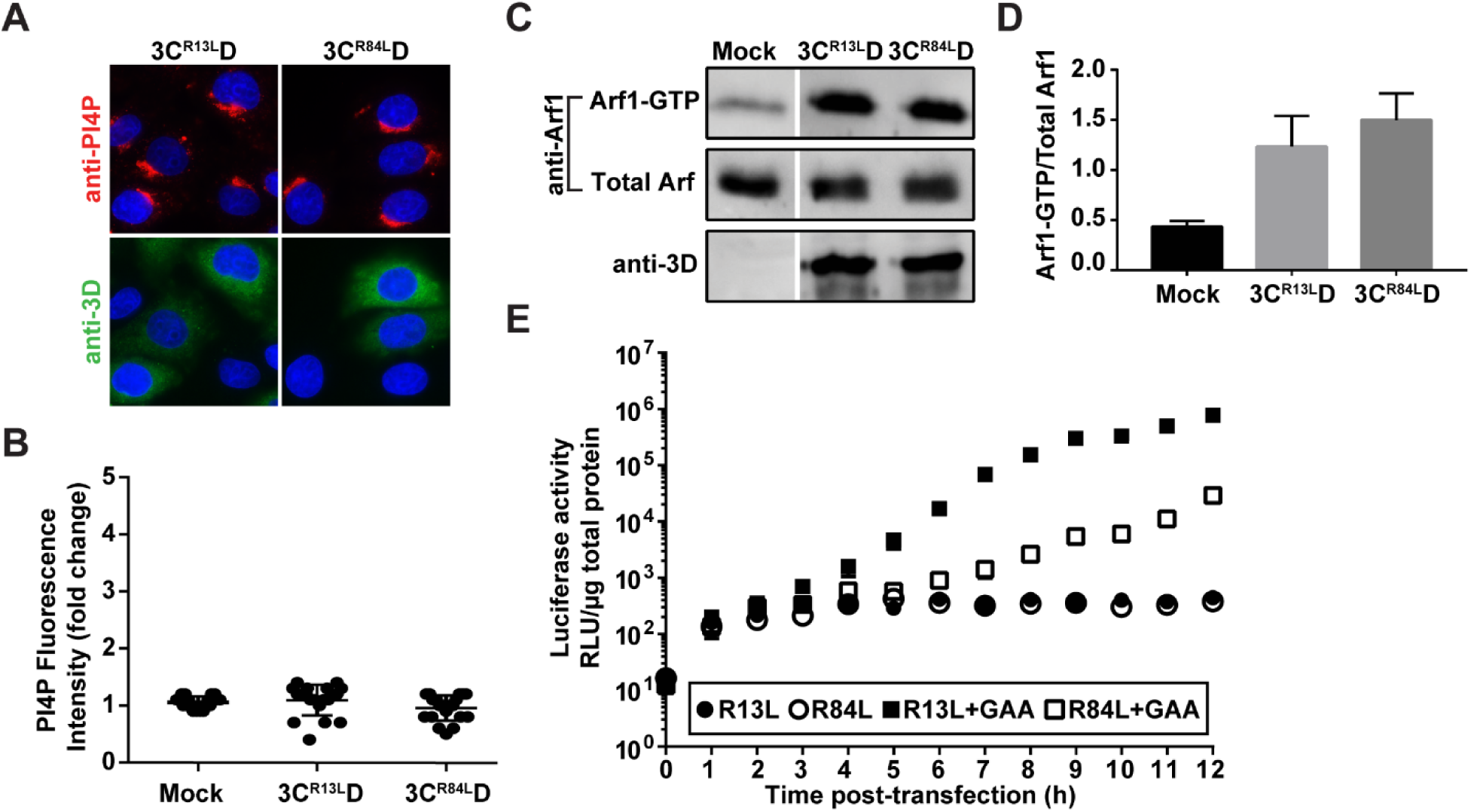
Additional 3CD variants with substitutions in the 3C domain exhibit defects to PI4P induction caused by block at a step post-Arf1 activation. **A** The indicated 3CD derivatives were expressed individually in HeLa cells and immunostained for the presence of PI4P (red) and 3CD (green). The nucleus was stained with DAPI (blue). Neither 3CD derivative altered the level of PI4P or its localization. **(B)** Quantification of PI4P intensity per cell (n=20) was performed as described in the legend to **Fig. 1C**. The average of the normalized values were: 1.06 ± 0.02 (SEM) in mock-transfected cells; 1.09 ± 0.06 (SEM) in 3C^R13L^D-transfected cells; 0.95 ± 0.05 (SEM) in 3C^R84L^D-transfected cells. The level of PI4P induction observed in 3CD mutant-transfected cells was not significant when compared to mock-transfected cells based on a Student’s t-test. **(C)** Activation of Arf1 by the 3CD derivatives was determined as described in the legend to **Fig. 3A**. Both derivatives induced activation of Arf1. **(D)** The magnitude of Arf1 activation was determined as described in the legend to **Fig. 3B**. The average of the values for the quotients (n=3) were: 0.43 ± 0.03 (SEM) in mock-transfected cells; 1.23 ± 0.18 (SEM) in 3C^R13L^D-transfected cells; 1.50 ± 0.15 (SEM) in 3C^R84L^D-transfected cells. For mock vs 3C^R13L^D, mock vs 3C^R84L^D and 3C^R13L^D vs 3C^R84L^D, a Student’s t-test yielded P values of 0.0112, 0.0024 and 0.3169, respectively. **(E)** Complementation of 3C^R13L^D-or 3C^R84L^D-expressing subgenomic replicons by 3CD-GAA. HeLa cells were transfected with replicon RNA indicated in the key. GAA refers to a replicon expressing a catalytically inactive 3D-encoded polymerase; the corresponding 3CD-GAA should function normally in PI4P induction. Luciferase activity was measured every hour post-transfection as indicated.

**Figure S3.**
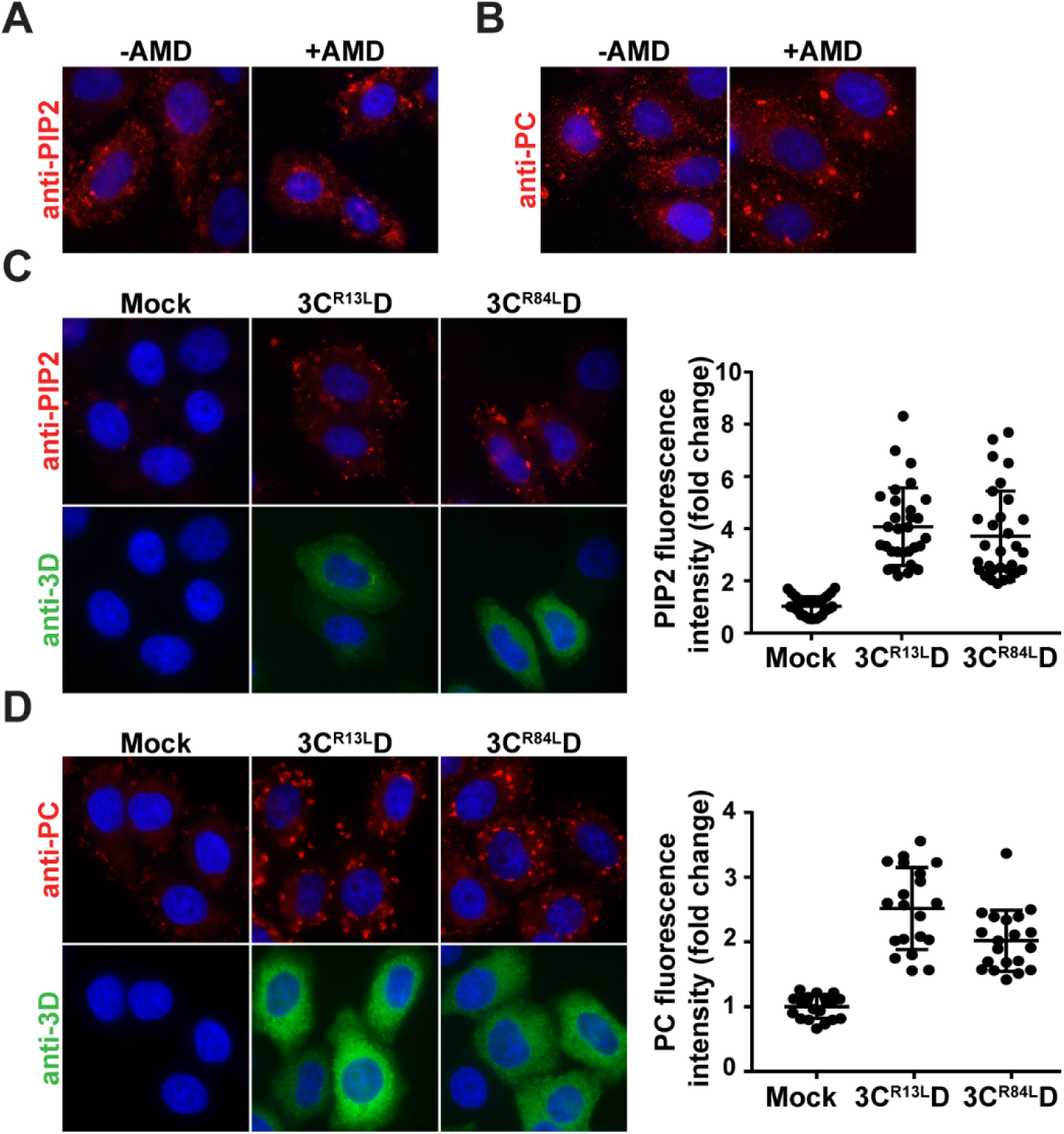
Induction of PIP2 and PC biosynthesis by 3CD is not sensitive to actinomycin D and does require induction of PI4P biosynthesis. **A** Effect of actinomycin D (AMD) on 3CD-mediated induction of PIP2 **(A)** and PC **(B)**. HeLa cells were maintained in the absence or presence of AMD (5 µg/mL) prior to transfection with 3CD mRNA. Cells were immunostained for PIP2 or PC (red) and 3CD (green); the nucleus was stained with DAPI (blue). The presence of AMD did not interfere with induction of PIP2 or PC. **(C)** The indicated 3CD derivatives were expressed individually in HeLa cells and immunostained for the presence of PIP2 (red) and 3CD (green). The nucleus was stained with DAPI (blue). Neither 3CD derivative impaired induction of PIP2. Quantification of PIP2 intensity per cell (n=30) was performed as described in the legend to **Fig. 1C**. The average of the normalized values were: 1.03 ± 0.07 (SEM) in mock-transfected cells; 4.71 ± 0.27 (SEM) in 3C^R13L^D-transfected cells; 4.08 ± 0.27 (SEM) in 3C^R84L^D-transfected cells. The level of PIP2 induction observed in 3CD mutant-transfected cells was not significantly different when compared to WT 3CD-transfected cells based on a Student’s t-test. **(D)** The indicated 3CD derivatives were expressed individually in HeLa cells and immunostained for the presence of PC (red) and 3CD (green). The nucleus was stained with DAPI (blue). Neither 3CD derivative impaired induction of PC. Quantification of PC intensity per cell (n=20) was performed as described in the legend to **Fig. 1C**. The average of the normalized values were: 1.00 ± 0.04 (SEM) in mock-transfected cells; 2.52 ± 0.14 (SEM) in 3C^R13L^D-transfected cells; 2.02 ± 0.11 (SEM) in 3C^R84L^D-transfected cells. The level of PC induction observed in 3CD mutant-transfected cells was not significantly different when compared to WT 3CD-transfected cells based on a Student’s t-test.

**Table S1.**
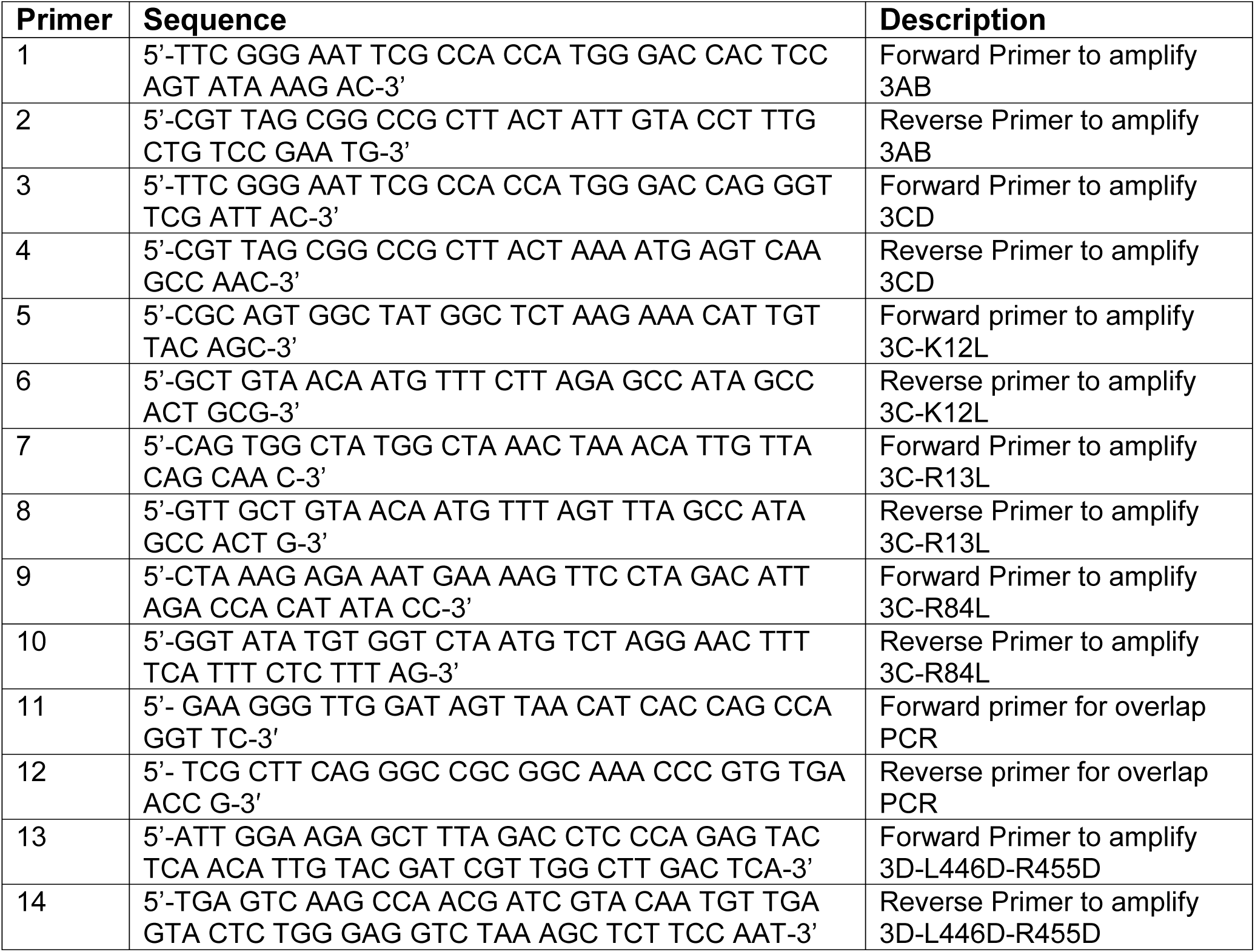
Oligonucleotides used in this study.

**Table S2.** Statistical analysis of all experimental data.

| Fig No. | Samples | Mean ± SEM | p-value | n |
| --- | --- | --- | --- | --- |
| Fig 1 | Mock | $1 \pm 0.04021$ | | 30 |
| | WT | $4.983 \pm 0.4259$ | | 30 |
|  | Mock vs. WT |  | <0.0001 |  |
| Fig 2 | WT-PIK93 | $1.841e^8 \pm 1.188e^7$ | | 25 |
| | WT+PIK93 | $7.192e^7 \pm 5.179e^6$ | | 25 |
|  | WT-PIK93 vs. WT+PIK93 |  | <0.0001 |  |
| Fig 3 | Mock | $0.4333 \pm 0.03333$ | | 3 |
| | WT | $1.767 \pm 0.03333$ | | 3 |
|  | Mock vs. WT |  | <0.0001 |  |
| Fig 4 (B) | Mock | $1 \pm 0.03748$ | | 30 |
| | 3CD<sup>m</sup> | $1.064 \pm 0.04205$ | | 30 |
| | 3C<sup>m</sup>D | $1.100 \pm 0.04400$ | | 30 |
| | 3CD<sup>m</sup>+3C<sup>m</sup>D | $3.083 \pm 0.1339$ | | 30 |
|  | Mock vs. 3CD<sup>m</sup> |  | 0.2602 |  |
|  | Mock vs. 3C<sup>m</sup>D |  | 0.089 |  |
|  | Mock vs. 3CD<sup>m</sup>+ 3C<sup>m</sup>D |  | <0.0001 |  |
|  | 3CD<sup>m</sup> vs. 3C<sup>m</sup>D |  | 0.5574 |  |
|  | 3CD<sup>m</sup> vs. 3CD<sup>m</sup>+ 3C<sup>m</sup>D |  | < 0.0001 |  |
|  | 3C<sup>m</sup>D vs. 3CD<sup>m</sup>+ 3C<sup>m</sup>D |  | < 0.0001 |  |
| Fig 4 (D) | Mock | $0.4246 \pm 0.01855$ | | 3 |
| | 3CD<sup>m</sup> | $0.3667 \pm 0.06667$ | | 3 |
| | 3C<sup>m</sup>D | $1.267 \pm 0.08819$ | | 3 |
|  | Mock vs. 3CD<sup>m</sup> |  | 0.4498 |  |
|  | Mock vs. 3C<sup>m</sup>D |  | 0.0007 |  |
|  | 3CD<sup>m</sup> vs. 3C<sup>m</sup>D |  | 0.0012 |  |
| Fig 5 (B) | WT | $170 \pm 10$ | | 3 |
| | 3CD<sup>m</sup> | $330 \pm 30$ | | 3 |
| | 3C<sup>m</sup>D | $80 \pm 10$ | | 3 |
|  | WT vs 3CD<sup>m</sup> |  | 0.0072 |  |
|  | WT vs 3C<sup>m</sup>D |  | 0.0031 |  |
|  | 3CD<sup>m</sup> vs 3C<sup>m</sup>D |  | 0.0014 |  |
| Fig 6 (A) | Mock | $1 \pm 0.04509$ | | 30 |
| | Infection | $6.624 \pm 0.3671$ | | 30 |
|  | Mock vs. Infection |  | <0.0001 |  |
| Fig 6 (B) | Mock | $1 \pm 0.04214$ | | 30 |
| | Infection | $3.033 \pm 0.1509$ | | 30 |
|  | Mock vs. Infection |  | <0.0001 |  |
| Fig 6 (C) | Mock | $1.000 \pm 0.06885$ | | 30 |
| | WT | $4.368 \pm 0.2510$ | | 30 |
| | 3CD<sup>m</sup> | $3.209 \pm 0.1580$ | | 30 |
| | 3C<sup>m</sup>D | $3.818 \pm 0.2512$ | | 30 |
|  | Mock vs. WT |  | <0.0001 |  |
|  | Mock vs. 3CD<sup>m</sup> |  | <0.0001 |  |
|  | Mock vs. 3C<sup>m</sup>D |  | <0.0001 |  |
|  | WT vs. 3CD<sup>m</sup> |  | 0.0002 |  |
|  | WT vs. 3C<sup>m</sup>D |  | 0.1264 |  |
|  | 3CD<sup>m</sup> vs. 3C<sup>m</sup>D |  | 0.0447 |  |
| Fig 6 (D) | Mock | $1 \pm 0.02185$ | | 45 |
| | WT | $2.18 \pm 0.07145$ | | 45 |
| | 3CD<sup>m</sup> | $1.887 \pm 0.06902$ | | 45 |
| | 3C<sup>m</sup>D | $2.057 \pm 0.1021$ | | 28 |
|  | Mock vs. WT |  | <0.0001 |  |
|  | Mock vs. 3CD<sup>m</sup> |  | <0.0001 |  |
|  | Mock vs. 3C<sup>m</sup>D |  | <0.0001 |  |
|  | WT vs. 3CD<sup>m</sup> |  | 0.0041 |  |
|  | WT vs. 3C<sup>m</sup>D |  | 0.3129 |  |
|  | 3CD<sup>m</sup> vs. 3C<sup>m</sup>D |  | 0.1583 |  |
| Fig S1 | Mock | $1.275 \pm 0.1436$ | | 4 |
| | Infection | $8.375 \pm 0.8087$ | | 4 |
|  | Mock vs. Infection |  | 0.0001 |  |
| Fig S2 (B) | Mock | $1.055 \pm 0.02348$ | | 20 |
| | 3C<sup>R13L</sup>D | $1.095 \pm 0.06003$ | | 20 |
| | 3C<sup>R84L</sup>D | $0.9550 \pm 0.04946$ | | 20 |
|  | Mock vs. 3C<sup>R13L</sup>D |  | 0.5386 |  |
|  | Mock vs. 3C<sup>R84L</sup>D |  | 0.0756 |  |
|  | 3C<sup>R13L</sup>D vs. 3C<sup>R84L</sup>D |  | 0.0798 |  |
| Fig S2 (D) | Mock | $0.4333 \pm 0.03333$ | | 3 |
| | 3C<sup>R13L</sup>D | $1.233 \pm 0.1764$ | | 3 |
| | 3C<sup>R84L</sup>D | $1.5 \pm 0.1528$ | | 3 |
|  | Mock vs. 3C<sup>R13L</sup>D |  | 0.0112 |  |
|  | Mock vs. 3C<sup>R84L</sup>D |  | 0.0024 |  |
|  | 3C<sup>R13L</sup>D vs. 3C<sup>R84L</sup>D |  | 0.3169 |  |
| Fig S3 (C) | Mock | $1.033 \pm 0.0659$ | | 30 |
| | WT | $4.711 \pm 0.2707$ | | 30 |
|  | 3C<sup>R13L</sup>D | 4.079 ± 0.2719 |  | 30 |
|  | 3C<sup>R84L</sup>D | 3.714 ± 0.314 |  | 30 |
|  | Mock vs. WT |  | <0.0001 |  |
|  | Mock vs. 3C<sup>R13L</sup>D |  | <0.0001 |  |
|  | Mock vs. 3C<sup>R84L</sup>D |  | <0.0001 |  |
|  | WT vs. 3C<sup>R13L</sup>D |  | 0.1053 |  |
|  | WT vs. 3C<sup>R84L</sup>D |  | 0.0194 |  |
|  | 3C<sup>R13L</sup>D vs. 3C<sup>R84L</sup>D |  | 0.3827 |  |
| Fig S3 (D) | Mock | 1 ± 0.04008 |  | 20 |
|  | WT | 2.389 ± 0.1503 |  | 20 |
|  | 3C<sup>R13L</sup>D | 2.517 ± 0.1417 |  | 20 |
|  | 3C<sup>R84L</sup>D | 2.021 ± 0.1055 |  | 20 |
|  | Mock vs. WT |  | <0.0001 |  |
|  | Mock vs. 3C<sup>R13L</sup>D |  | <0.0001 |  |
|  | Mock vs. 3C<sup>R84L</sup>D |  | <0.0001 |  |
|  | WT vs. 3C<sup>R13L</sup>D |  | 0.5371 |  |
|  | WT vs. 3C<sup>R84L</sup>D |  | 0.0523 |  |
|  | 3C<sup>R13L</sup>D vs. 3C<sup>R84L</sup>D |  | 0.0078 |  |

## REFERENCES

1. Feng Q, Langereis MA, van Kuppeveld FJ (2014) Induction and suppression of innate antiviral responses by picornaviruses. Cytokine Growth Factor Rev 25: 577–585.

2. Lei X, Xiao X, Wang J (2016) Innate Immunity Evasion by Enteroviruses: Insights into Virus-Host Interaction. Viruses 8.

3. Lu J, Yi L, Ke C, Zhang Y, Liu R, et al. (2015) The interaction between human enteroviruses and type I IFN signaling pathway. Crit Rev Microbiol 41: 201–207.

4. Pathinayake PS, Hsu AC, Wark PA (2015) Innate Immunity and Immune Evasion by Enterovirus 71. Viruses 7: 6613–6630.

5. Scutigliani EM, Kikkert M (2017) Interaction of the innate immune system with positive-strand RNA virus replication organelles. Cytokine Growth Factor Rev 37: 17–27.

6. Melia CE, van der Schaar HM, Lyoo H, Limpens R, Feng Q, et al. (2017) Escaping Host Factor PI4KB Inhibition: Enterovirus Genomic RNA Replication in the Absence of Replication Organelles. Cell Rep 21: 587–599.

7. den Boon JA, Ahlquist P (2010) Organelle-like membrane compartmentalization of positive-strand RNA virus replication factories. Annu Rev Microbiol 64: 241–256.

8. den Boon JA, Diaz A, Ahlquist P (2010) Cytoplasmic viral replication complexes. Cell Host Microbe 8: 77–85.

9. Paul D, Bartenschlager R (2015) Flaviviridae Replication Organelles: Oh, What a Tangled Web We Weave. Annu Rev Virol 2: 289–310.

10. Jose J, Snyder JE, Kuhn RJ (2009) A structural and functional perspective of alphavirus replication and assembly. Future Microbiol 4: 837–856.

11. Belov GA, Nair V, Hansen BT, Hoyt FH, Fischer ER, et al. (2012) Complex dynamic development of poliovirus membranous replication complexes. J Virol 86: 302–312.

12. Chatel-Chaix L, Bartenschlager R (2014) Dengue virus-and hepatitis C virus-induced replication and assembly compartments: the enemy inside—caught in the web. J Virol 88: 5907–5911.

13. Meyers NL, Fontaine KA, Kumar GR, Ott M (2016) Entangled in a membranous web: ER and lipid droplet reorganization during hepatitis C virus infection. Curr Opin Cell Biol 41: 117–124.

14. Richards AL, Soares-Martins JA, Riddell GT, Jackson WT (2014) Generation of unique poliovirus RNA replication organelles. MBio 5: e00833-00813.

15. Ahlquist P, Noueiry AO, Lee WM, Kushner DB, Dye BT (2003) Host factors in positive-strand RNA virus genome replication. J Virol 77: 8181–8186.

16. Hsu NY, Ilnytska O, Belov G, Santiana M, Chen YH, et al. (2010) Viral reorganization of the secretory pathway generates distinct organelles for RNA replication. Cell 141: 799–811.

17. Balla T (2013) Phosphoinositides: tiny lipids with giant impact on cell regulation. Physiol Rev 93: 1019–1137.

18. Weixel KM, Blumental-Perry A, Watkins SC, Aridor M, Weisz OA (2005) Distinct Golgi populations of phosphatidylinositol 4-phosphate regulated by phosphatidylinositol 4-kinases. J Biol Chem 280: 10501–10508.

19. Dornan GL, McPhail JA, Burke JE (2016) Type III phosphatidylinositol 4 kinases: structure, function, regulation, signalling and involvement in disease. Biochem Soc Trans 44: 260–266.

20. Donaldson JG, Honda A, Weigert R (2005) Multiple activities for Arf1 at the Golgi complex. Biochim Biophys Acta 1744: 364–373.

21. Kaczmarek B, Verbavatz JM, Jackson CL (2017) GBF1 and Arf1 function in vesicular trafficking, lipid homeostasis and organelle dynamics. Biol Cell.

22. Altan-Bonnet N, Balla T (2012) Phosphatidylinositol 4-kinases: hostages harnessed to build panviral replication platforms. Trends Biochem Sci 37: 293–302.

23. Zhang L, Hong Z, Lin W, Shao RX, Goto K, et al. (2012) ARF1 and GBF1 generate a PI4P-enriched environment supportive of hepatitis C virus replication. PLoS One 7: e32135.

24. Wessels E, Duijsings D, Niu TK, Neumann S, Oorschot VM, et al. (2006) A viral protein that blocks Arf1-mediated COP-I assembly by inhibiting the guanine nucleotide exchange factor GBF1. Dev Cell 11: 191–201.

25. Xiao X, Lei X, Zhang Z, Ma Y, Qi J, et al. (2017) Enterovirus 3A facilitates viral replication by promoting PI4KB-ACBD3 interaction. J Virol.

26. Dorobantu CM, Ford-Siltz LA, Sittig SP, Lanke KH, Belov GA, et al. (2015) GBF1-and ACBD3-independent recruitment of PI4KIIIbeta to replication sites by rhinovirus 3A proteins. J Virol 89: 1913–1918.

27. Dorobantu CM, van der Schaar HM, Ford LA, Strating JR, Ulferts R, et al. (2014) Recruitment of PI4KIIIbeta to coxsackievirus B3 replication organelles is independent of ACBD3, GBF1, and Arf1. J Virol 88: 2725–2736.

28. Greninger AL, Knudsen GM, Betegon M, Burlingame AL, Derisi JL (2012) The 3A protein from multiple picornaviruses utilizes the golgi adaptor protein ACBD3 to recruit PI4KIIIbeta. J Virol 86: 3605–3616.

29. Sasaki J, Ishikawa K, Arita M, Taniguchi K (2012) ACBD3-mediated recruitment of PI4KB to picornavirus RNA replication sites. EMBO J 31: 754–766.

30. Pathak HB, Oh HS, Goodfellow IG, Arnold JJ, Cameron CE (2008) Picornavirus genome replication: roles of precursor proteins and rate-limiting steps in oriIdependent VPg uridylylation. J Biol Chem 283: 30677–30688.

31. Oh HS, Pathak HB, Goodfellow IG, Arnold JJ, Cameron CE (2009) Insight into poliovirus genome replication and encapsidation obtained from studies of 3B-3C cleavage site mutants. J Virol 83: 9370–9387.

32. Sandoval IV, Carrasco L (1997) Poliovirus infection and expression of the poliovirus protein 2B provoke the disassembly of the Golgi complex, the organelle target for the antipoliovirus drug Ro-090179. J Virol 71: 4679–4693.

33. Doedens JR, Giddings TH, Jr., Kirkegaard K (1997) Inhibition of endoplasmic reticulum-to-Golgi traffic by poliovirus protein 3A: genetic and ultrastructural analysis. J Virol 71: 9054–9064.

34. Doedens JR, Kirkegaard K (1995) Inhibition of cellular protein secretion by poliovirus proteins 2B and 3A. EMBO J 14: 894–907.

35. Sharma R, Raychaudhuri S, Dasgupta A (2004) Nuclear entry of poliovirus protease-polymerase precursor 3CD: implications for host cell transcription shutoff. Virology 320: 195–205.

36. Bensaude O (2011) Inhibiting eukaryotic transcription: Which compound to choose? How to evaluate its activity? Transcription 2: 103–108.

37. Luna L, Rolseth V, Hildrestrand GA, Otterlei M, Dantzer F, et al. (2005) Dynamic relocalization of hOGG1 during the cell cycle is disrupted in cells harbouring the hOGG1-Cys326 polymorphic variant. Nucleic Acids Res 33: 1813–1824.

38. Saenz JB, Sun WJ, Chang JW, Li J, Bursulaya B, et al. (2009) Golgicide A reveals essential roles for GBF1 in Golgi assembly and function. Nat Chem Biol 5: 157–165.

39. Toth B, Balla A, Ma H, Knight ZA, Shokat KM, et al. (2006) Phosphatidylinositol 4kinase IIIbeta regulates the transport of ceramide between the endoplasmic reticulum and Golgi. J Biol Chem 281: 36369–36377.

40. Santy LC, Casanova JE (2001) Activation of ARF6 by ARNO stimulates epithelial cell migration through downstream activation of both Rac1 and phospholipase D. J Cell Biol 154: 599–610.

41. Spear A, Ogram SA, Morasco BJ, Smerage LE, Flanegan JB (2015) Viral precursor protein P3 and its processed products perform discrete and essential functions in the poliovirus RNA replication complex. Virology 485: 492–501.

42. Towner JS, Mazanet MM, Semler BL (1998) Rescue of defective poliovirus RNA replication by 3AB-containing precursor polyproteins. J Virol 72: 7191–7200.

43. Pathak HB, Ghosh SK, Roberts AW, Sharma SD, Yoder JD, et al. (2002) Structure-function relationships of the RNA-dependent RNA polymerase from poliovirus (3Dpol). A surface of the primary oligomerization domain functions in capsid precursor processing and VPg uridylylation. J Biol Chem 277: 31551–31562.

44. Shengjuler D, Sun S, Cremer PS, Cameron CE (2017) PIP-on-a-chip: A Label-free Study of Protein-phosphoinositide Interactions. J Vis Exp.

45. Vance DE, Trip EM, Paddon HB (1980) Poliovirus increases phosphatidylcholine biosynthesis in HeLa cells by stimulation of the rate-limiting reaction catalyzed by CTP: phosphocholine cytidylyltransferase. J Biol Chem 255: 1064–1069.

46. Zhang J, Zhang Z, Chukkapalli V, Nchoutmboube JA, Li J, et al. (2016) Positive-strand RNA viruses stimulate host phosphatidylcholine synthesis at viral replication sites. Proc Natl Acad Sci U S A 113: E1064–1073.

47. Belov GA, Habbersett C, Franco D, Ehrenfeld E (2007) Activation of cellular Arf GTPases by poliovirus protein 3CD correlates with virus replication. J Virol 81: 9259–9267.

48. Boura E, Nencka R (2015) Phosphatidylinositol 4-kinases: Function, structure, and inhibition. Exp Cell Res 337: 136–145.

49. Arita M (2016) Mechanism of Poliovirus Resistance to Host Phosphatidylinositol-4 Kinase III beta Inhibitor. ACS Infect Dis 2: 140–148.

50. van der Schaar HM, van der Linden L, Lanke KH, Strating JR, Purstinger G, et al. (2012) Coxsackievirus mutants that can bypass host factor PI4KIIIbeta and the need for high levels of PI4P lipids for replication. Cell Res 22: 1576–1592.

51. Tan X, Thapa N, Choi S, Anderson RA (2015) Emerging roles of PtdIns(4,5)P2-beyond the plasma membrane. J Cell Sci 128: 4047–4056.

52. Tan X, Thapa N, Liao Y, Choi S, Anderson RA (2016) PtdIns(4,5)P2 signaling regulates ATG14 and autophagy. Proc Natl Acad Sci U S A 113: 10896–10901.

53. Bird SW, Maynard ND, Covert MW, Kirkegaard K (2014) Nonlytic viral spread enhanced by autophagy components. Proc Natl Acad Sci U S A 111: 13081-13086.

54. Jackson WT, Giddings TH, Jr., Taylor MP, Mulinyawe S, Rabinovitch M, et al. (2005) Subversion of cellular autophagosomal machinery by RNA viruses. PLoS Biol 3: e156.

55. Kirkegaard K, Jackson WT (2005) Topology of double-membraned vesicles and the opportunity for non-lytic release of cytoplasm. Autophagy 1: 182–184.

56. Klein KA, Jackson WT (2011) Picornavirus subversion of the autophagy pathway. Viruses 3: 1549–1561.

57. Lai JK, Sam IC, Chan YF (2016) The Autophagic Machinery in Enterovirus Infection. Viruses 8.

58. Richards AL, Jackson WT (2013) How positive-strand RNA viruses benefit from autophagosome maturation. J Virol 87: 9966–9972.

59. Schlegel A, Giddings TH, Jr., Ladinsky MS, Kirkegaard K (1996) Cellular origin and ultrastructure of membranes induced during poliovirus infection. J Virol 70: 65766588.

60. Suhy DA, Giddings TH, Jr., Kirkegaard K (2000) Remodeling the endoplasmic reticulum by poliovirus infection and by individual viral proteins: an autophagy-like origin for virus-induced vesicles. J Virol 74: 8953–8965.

61. Taylor MP, Kirkegaard K (2007) Modification of cellular autophagy protein LC3 by poliovirus. J Virol 81: 12543–12553.

62. Taylor MP, Kirkegaard K (2008) Potential subversion of autophagosomal pathway by picornaviruses. Autophagy 4: 286–289.

63. Chen YH, Du W, Hagemeijer MC, Takvorian PM, Pau C, et al. (2015) Phosphatidylserine vesicles enable efficient en bloc transmission of enteroviruses. Cell 160: 619–630.

64. van den Bout I, Divecha N (2009) PIP5K-driven PtdIns(4,5)P2 synthesis: regulation and cellular functions. J Cell Sci 122: 3837–3850.

65. Gustin KE, Sarnow P (2001) Effects of poliovirus infection on nucleo-cytoplasmic trafficking and nuclear pore complex composition. EMBO J 20: 240–249.

66. Cho NJ, Lee C, Pang PS, Pham EA, Fram B, et al. (2015) Phosphatidylinositol 4,5bisphosphate is an HCV NS5A ligand and mediates replication of the viral genome. Gastroenterology 148: 616–625.

67. Saad JS, Miller J, Tai J, Kim A, Ghanam RH, et al. (2006) Structural basis for targeting HIV-1 Gag proteins to the plasma membrane for virus assembly. Proc Natl Acad Sci U S A 103: 11364–11369.

68. Johnson KA, Taghon GJ, Scott JL, Stahelin RV (2016) The Ebola Virus matrix protein, VP40, requires phosphatidylinositol 4,5-bisphosphate (PI(4,5)P2) for extensive oligomerization at the plasma membrane and viral egress. Sci Rep 6: 19125.

69. Kakisaka M, Yamada K, Yamaji-Hasegawa A, Kobayashi T, Aida Y (2016) Intrinsically disordered region of influenza A NP regulates viral genome packaging via interactions with viral RNA and host PI(4,5)P2. Virology 496: 116–126.

70. Moustafa IM, Gohara DW, Uchida A, Yennawar N, Cameron CE (2015) Conformational Ensemble of the Poliovirus 3CD Precursor Observed by MD Simulations and Confirmed by SAXS: A Strategy to Expand the Viral Proteome? Viruses 7: 5962–5986.

71. Arnold JJ, Bernal A, Uche U, Sterner DE, Butt TR, et al. (2006) Small ubiquitin-like modifying protein isopeptidase assay based on poliovirus RNA polymerase activity. Anal Biochem 350: 214–221.

